# Autophagy acts as a brake on obesity-related fibrosis by controlling purine nucleoside signalling

**DOI:** 10.1101/2024.09.17.613382

**Authors:** Klara Piletic, Amir H. Kayvanjoo, Felix Clemens Richter, Mariana Borsa, Ana V. Lechuga-Vieco, Oliver Popp, Sacha Grenet, Jacky Ka Long Ko, Kristina Zec, Maria Kyriazi, Lada Koneva, Stephen Sansom, Philipp Mertins, Fiona Powrie, Ghada Alsaleh, Anna Katharina Simon

## Abstract

A hallmark of obesity is a pathological expansion of white adipose tissue (WAT), accompanied by marked tissue dysfunction and fibrosis. Autophagy promotes adipocyte differentiation and lipid homeostasis, but its role in obese adipocytes and adipose tissue dysfunction remains incompletely understood. Here, we demonstrate that autophagy is a key tissue-specific regulator of WAT remodelling in diet-induced obesity. Importantly, loss of adipocyte autophagy substantially exacerbates pericellular fibrosis in visceral WAT. Change in WAT architecture correlates with increased infiltration of macrophages with tissue-reparative, fibrotic features. We uncover that autophagy regulates purine nucleoside metabolism in obese adipocytes, preventing excessive release of the purine catabolites xanthine and hypoxanthine. Purines signal cell-extrinsically for fibrosis by driving macrophage polarisation towards a tissue reparative phenotype. Our findings reveal a novel role for adipocyte autophagy in regulating tissue purine nucleoside metabolism, thereby limiting obesity-associated fibrosis and maintaining the functional integrity of visceral WAT. Purine signals may serve as a critical balance checkpoint and therapeutic target in fibrotic diseases.

## INTRODUCTION

Excess weight and obesity represent a major global health and socioeconomic burden (González-Muniesa et al., 2017). Obesity pathogenesis is characterized by a marked increase in white adipose tissue (WAT) mass, predominantly in subcutaneous and visceral locations, with the latter being more detrimental in obesity pathophysiology (Rosen and Spiegelman, 2014). Excess adiposity is considered a major risk factor for metabolic complications, including type II diabetes mellitus and fatty liver disease (Sakers et al., 2022). While traditionally viewed as a highly specialized tissue for energy storage and mobilization, adipose tissue is now recognized as a dynamic endocrine and paracrine organ (Ghaben and Scherer, 2019). During excess nutrient availability, WAT mass increases through cell growth (hypertrophy) and number (hyperplasia) of adipocytes, which shift their metabolism to meet the energetic demands of the organism (Chouchani and Kajimura, 2019). The obesity-associated metabolic shift predominantly includes core lipid and glucose metabolism to support energy storage and mobilisation, and these processes are tightly linked to functional mitochondria (Morigny et al., 2021). However, adipocyte metabolism and obesity-related metabolic rewiring beyond these pathways remain poorly understood. Adipose tissue architectural changes are supported by a dynamic remodelling of extracellular matrix (ECM). Rapid and chronic expansion leads to hypoxia, chronic low-grade inflammation, and fibrosis, rendering WAT inflexible and dysfunctional (Gliniak et al., 2023). Besides obesity-induced changes in adipocytes, adipose tissue dysfunction is also characterized by an accumulation of adipose tissue macrophages (ATMs) (Matz et al., 2023). ATMs create an inflammatory milieu by releasing inflammatory cytokines and support adipose tissue fibrogenesis through ECM turnover and fibroblast stimulation (Marcelin et al., 2022). While the exact sequence of these processes is still unclear (Reggio et al., 2013, Chait and den Hartigh, 2020), their detrimental impact on adipose tissue is indisputable.

Autophagy is a fundamental process for the regulation of cellular metabolism and energy homeostasis (Kaur and Debnath, 2015). Through a highly dynamic regulation of cellular recycling and degradation, autophagy controls metabolic adaptation, differentiation, homeostasis, and ultimately the overall function of cells and organs (Dikic and Elazar, 2018). Autophagy is activated by various cellular and environmental stress signals, including nutrient and energy deprivation, and oxidative stress (Klionsky et al., 2021). When initiated, it recycles organelles and macromolecules either as a quality control mechanism or to replenish energy and anabolic precursor pools. Through these processes, it can both rewire metabolic processes as well as supply nutrients, deeming it a master regulator of cellular metabolism (Rabinowitz and White, 2010, Deretic and Kroemer, 2022). Notably, autophagy can supply nutrients both in a cell-intrinsic as well as in a cell-extrinsic manner (Piletic et al., 2023).

Autophagy supports adipocyte differentiation and lipid homeostasis, as well as facilitates communication between adipose tissue and the liver (Singh et al., 2009, Zhang et al., 2009, Cai et al., 2018, Sakane et al., 2021), however, its function in obese adipocytes and adipose tissue dysfunction remains unclear and controversial (Clemente-Postigo et al., 2020, Soussi et al., 2016). Here we demonstrate that in obese conditions, adipocytes upregulate autophagy to support their metabolic and structural adaptation. Failure to meet their metabolic demands in the absence of autophagy leads to elevated purine nucleoside production and secretion. Xanthine and hypoxanthine-mediated adipocyte-macrophage crosstalk drives a tissue-reparative macrophage phenotype and ultimately leads to excessive pericellular adipose tissue fibrosis.

## RESULTS

### Autophagy is dysregulated in obesity, regulating WAT remodelling by limiting pericellular fibrosis

Adipocytes undergo significant structural, metabolic, and functional remodelling during obesity. To gain deeper insight into how obesity alters human WAT adipocytes, we analysed a recently published human WAT single-nuclei RNAseq (sn-RNAseq) atlas (Emont et al., 2022). Comparison of white adipocytes between lean and obese states revealed macroautophagy as one of the key dysregulated pathways, together with multiple well-studied pathways impacted by weight gain, including insulin signalling, lipid metabolism, and tissue repair (Fig 1A-B). Accordingly, we observed that in mice fed with high fat diet (HFD) to induce obesity, autophagy initially correlated positively with increased adiposity. This observation correlated with previous reports of autophagy upregulation during obesity (Clemente-Postigo et al., 2020, Jansen et al., 2012, Kosacka et al., 2015, Kovsan et al., 2011, Mizunoe et al., 2017, Nuñez et al., 2013, Öst et al., 2010). Prolonged HFD feeding, however, led to a significant downregulation of autophagic flux (Fig 1C). To investigate its pathophysiological role, we generated a mouse model with an inducible, adipocyte-specific deletion of *Atg7* (*Atg7^Ad^*) to circumvent defective adipogenesis (Richter et al., 2023). Following the induction of *Atg7* deletion in mature adipocytes (Fig S1A), the obesity-associated increase in autophagy flux was still abrogated in gonadal WAT (gWAT) adipose depot 16 weeks after the tamoxifen treatment (Fig S1B-C). Loss of autophagy in adipocytes had a profound impact on the tissue structure of obese *Atg7^Ad^* gWAT (Fig 1D). Obese *Atg7^Ad^* mice showed exacerbated pericellular fibrosis compared to littermate controls (Fig 1E-F), which correlated with an increase in obesity-induced gWAT autophagy in WT mice (Fig 1C). Pericellular fibrosis onset in gWAT developed during the initial body weight gain between six and nine weeks of high fat diet feeding (Fig 1E-F). In line with the aggravated ECM accumulation, several ECM components and enzymes, including *Col3a1*, *Fn1*, *Mmp14*, and *Timp1*, were strongly increased (Fig 1G). Taken together, these data point towards a critical role of obesity-induced autophagy in the control of ECM remodelling and tissue fibrosis.

**Figure 1:**
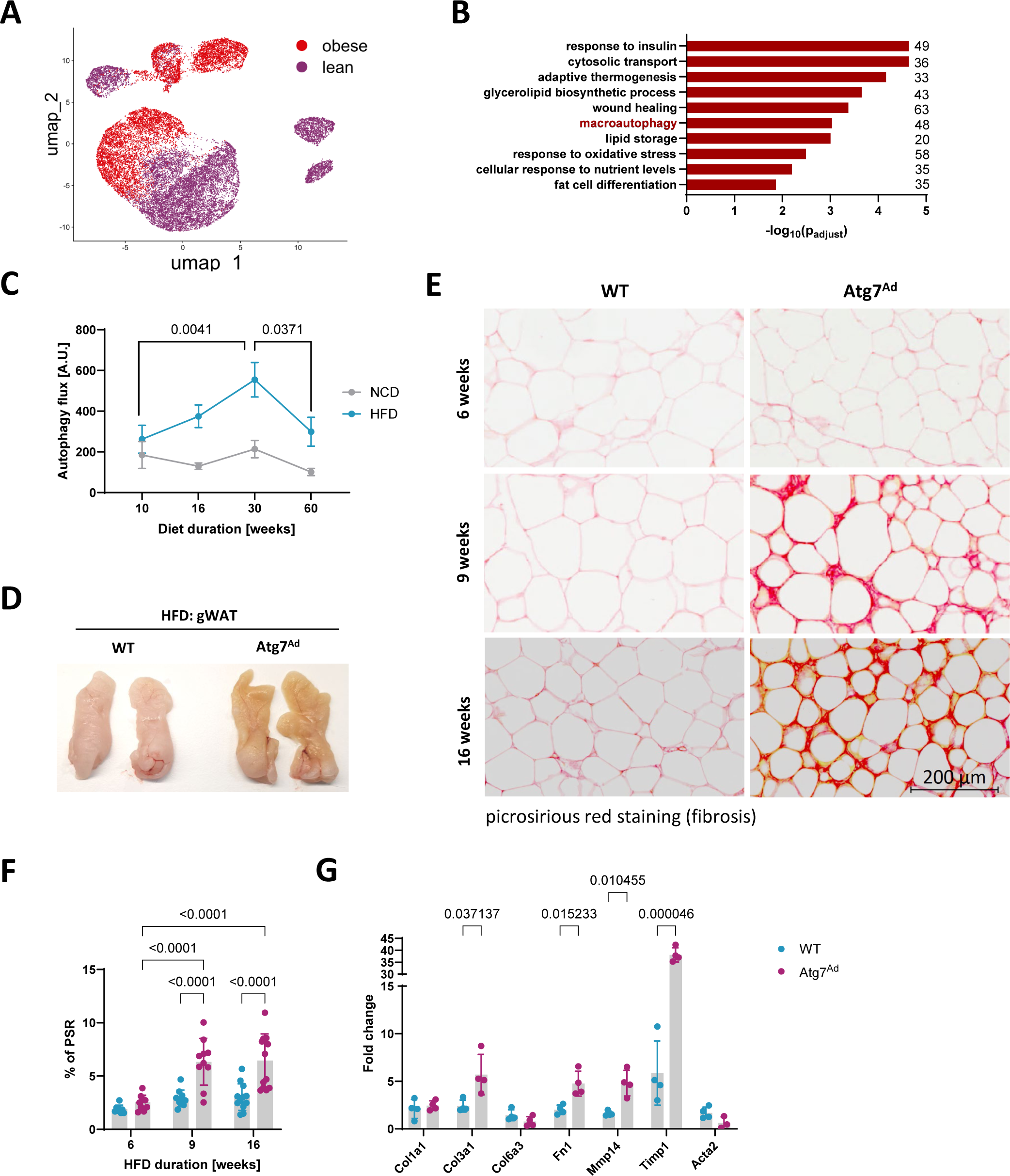
Obesity dysregulates autophagy and limits pericellular fibrosis in WAT. A) UMAP projection of human white adipocytes from lean (BMI < 30; 12822 adipocytes) and obese (BMI > 40; 9191 adipocytes) subjects. Single nucleus RNA-seq data has been obtained from a deposited dataset (GSE176171). B) Enrichment GO analysis of differentially regulated pathways in human adipocytes isolated from obese compared to lean WAT. The number of genes identified for each term is labelled. C) WT mice were fed a normal chow diet (NCD) or high fat diet (HFD) for 10, 30 or 60 weeks before autophagy flux in gonadal white adipose tissue (gWAT) was assessed as explained in Materials and Methods. Western blot analysis of autophagy flux was calculated as (LC3-II (Inh) – LC3-II (Veh)). n = 5 – 8 mice. Data are merged from 3 independent experiments. D) Photograph of gWAT fat pads of WT and *Atg7^Ad^* mice fed with high fat diet (HFD) for 16 weeks. Two representatives of 3 independent experiments. E) Picrosirius red staining (PSR), specifically staining collagen I and III, of gWAT depots harvested from HFD-fed WT and *Atg7^Ad^* mice after 6, 9, and 16 weeks of feeding. Representative images are shown. Scale bar, 200 µm. F) Quantification of picrosirius red positive area as a percentage of total area from (E). n = 5-10 mice. Data are merged from 3 independent experiments. G) Relative mRNA levels of ECM-related genes in gWAT after 16 weeks of HFD measured by qRT-PCR. n = 3-4 mice. Representative of 3 independent experiments. Data are presented as mean ± SEM (C) or mean ± SD (F-G). Dots represent individual biological replicates. Statistical analysis by two-way ANOVA with Tukey multi comparisons (C) or Fisher (F test or multiple unpaired t-test (G).

### Multi-OMICS analysis reveals a key role for autophagy in adipocyte metabolic adaptation and nucleotide homeostasis during obesity

We next asked whether a striking shift in fibrotic processes in obese *Atg7^Ad^* gWAT was due to an autophagy-mediated cell-intrinsic process. To explore the role of autophagy in adipocyte cellular remodelling, we conducted a multi-OMICS analysis of obese WT and *Atg7^Ad^* adipocytes (Fig 2). Proteomics analysis showed that classical autophagy receptors such as SQSTM1, which are typically degraded during autophagy, accumulated in autophagy-deficient adipocytes, confirming a lack of autophagy function (Fig S2A). The analysis highlighted impaired mitochondrial homeostasis in *Atg7^Ad^* adipocytes, evidenced by reduced expression of the electron transport chain subunit complexes I-V following autophagy ablation (Fig S2B). Furthermore, we found that the absence of autophagy moderately affected adipocyte viability, as indicated by adipocyte marker Perilipin-1 staining (Fig S2C-D). The impact of autophagy loss on viability, however, was considerably lower compared to the impact of HFD feeding and obesity. Treatment with the pan-caspase inhibitor Q-VD-OPh (Caserta et al., 2003) reduced adipocyte cell death (Fig S2E) suggesting cell death occurred via caspase 3-induced apoptosis. Unexpectedly a decrease in adipokine production and secretion, including leptin, adiponectin, and Dpp4 (Fig S2F-G) was revealed. Among the significantly enriched proteins (Fig 2A), loss of adipocyte autophagy significantly altered metabolic processes, particularly nucleotide and lipid metabolism, as well as responses to oxidative stress (Fig 2B-C and Fig S2A).

**Figure 2:**
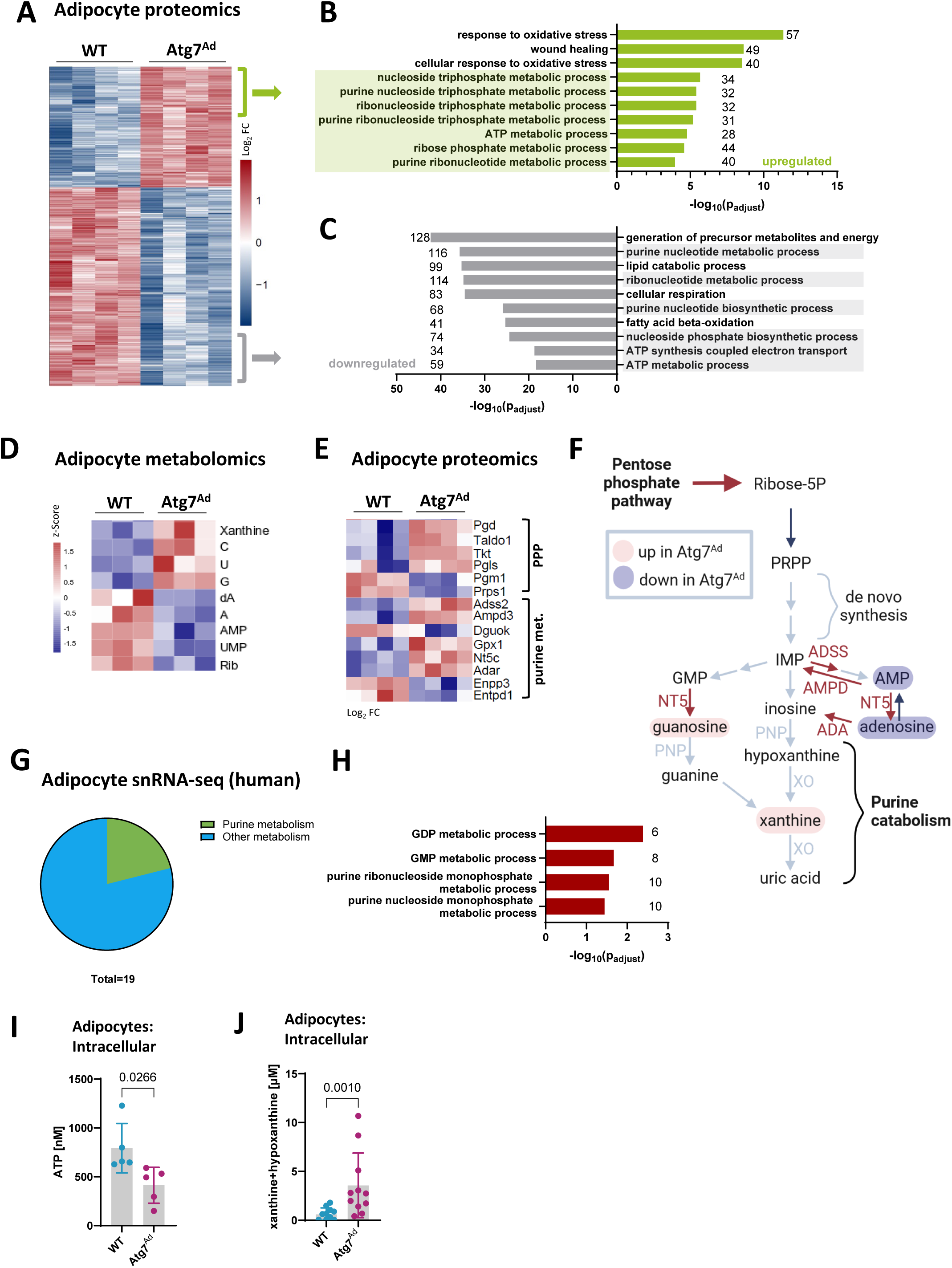
Autophagy controls adipocyte purine nucleoside metabolism. A-C) Hierarchical clustering of proteomics profiles of enriched proteins in adipocytes isolated from gWAT of WT and *Atg7^Ad^* mice fed with HFD for 16 weeks (1886 proteins identified with adjusted p-value ≤ 0.01). Colour-coding represents the log2 fold difference between WT and *Atg7^Ad^*mice (A). Enrichment GO analysis of upregulated (B) or downregulated (C) pathways in adipocytes isolated from *Atg7^Ad^* compared to WT gWAT. The number of genes identified for each term is labelled. n = 4 mice. D) Z-score heatmap of significantly (p < 0.05) abundant nucleotide and nucleoside metabolites in adipocytes. Metabolomics analysis was performed on adipocytes isolated from gWAT of WT and *Atg7^Ad^* mice following HFD feeding for 16 weeks. n = 3 mice. E) Log2 fold change heatmap of significantly differentially abundant proteins between WT and *Atg7^Ad^* involved in PPP (pentose phosphate pathway) and purine nucleoside metabolism in adipocytes as measured by proteomics analysis (as in A). F) Schematic summary of adipocyte proteome and metabolome changes upon loss of autophagy depicting simplified pentose phosphate and purine nucleoside metabolic pathways. Representative enzymes and metabolic products are colour-coded based on the fold change (red ∼ upregulated, blue ∼ downregulated). G) Pie chart representation of a total of 19 metabolic pathways identified in the enrichment GO analysis of differentially regulated pathways in human adipocyte snRNA-seq isolated from obese compared to lean WAT. Four out of 19 relate to purine nucleoside metabolism. H) Differentially regulated GO pathways related to purine nucleoside metabolism in human adipocytes isolated from obese and lean subjects. The number of genes identified for each term is labelled. I-J) Concentration of intracellular ATP (I) and xanthine and hypoxanthine (J) in gWAT adipocytes from WT and *Atg7^Ad^* mice fed with HFD for 16 weeks. n = 5-11 mice. Representative of 3 independent experiments (I) or merged from 3 independent experiments (J). Data are presented as mean ± SD. Dots represent individual biological replicates. Statistical analysis by unpaired t-test (I) or Mann-Whitney test (J). All heatmap values were scaled by row (protein/metabolite) using z-score.

In line with our proteomics data, we found that adipocyte autophagy loss in obese mice profoundly impacted their cellular metabolome (Fig 2D and S3A). We observed that loss of autophagy led to a global reduction in both essential and non-essential amino acids, alongside enzymes supporting the TCA cycle and fatty acid oxidation (FAO) in obese adipocytes assessed by metabolomics and proteomics, respectively (Fig S3D-F). Furthermore, we found key components for RNA synthesis, including nucleotides UMP, and AMP, adenosine, as well as ribose significantly reduced (Fig 2D).

Surprisingly, the loss of autophagy resulted in a pronounced upregulation of metabolites primarily associated with nucleoside metabolism (Fig 2D). We found a strong accumulation of purine and pyrimidine nucleosides in autophagy-deficient adipocytes, including guanosine, cytidine, uridine, as well as xanthine, a downstream product of purine catabolism (Fig 2D). The dysregulation in nucleoside metabolism was further emphasised by altered protein levels of several critical enzymes involved in the pentose phosphate pathway (PPP) and intracellular purine metabolism (Fig 2E). A prominent increase in PPP and purine metabolism enzymes was observed alongside elevated glycolytic enzymes that support this metabolic axis (Fig S3F). Notably, these profound changes in adipocyte metabolism revealed that autophagy plays a critical role in the maintenance of functional nucleotide pools in adipocytes during obesity (Fig 2F). Remarkably, when examining snRNA-seq data of human adipocytes, we found a similar pattern; out of 19 dysregulated metabolic pathways with obesity, more than one-fifth were related to purine metabolism (Fig 2G-H). This parallel between mouse and human data underscores the central role of purine nucleoside metabolism in adipocytes and its regulation by autophagy during obesity.

To validate the role of autophagy in regulating purine metabolism, we measured both upstream and downstream intermediate metabolites with enzymatic assays, including ATP, hypoxanthine, xanthine, and uric acid. Obese *Atg7^Ad^* adipocytes showed a significant reduction in the energy-rich purine nucleotide ATP (Fig 2I). In contrast, downstream intermediates of purine catabolism xanthine and hypoxanthine were significantly increased (Fig 2J). However, there was no difference in the end product of purine catabolism, uric acid, or in the enzymatic activity of xanthine oxidase (Fig S3B-C), suggesting that purine catabolism was not upregulated to generate excess uric acid. Taken together, autophagy is indispensable to maintaining balanced purine nucleoside metabolism in obese adipocytes. Its impairment shifts the metabolism towards the increased catabolic activity of the nucleoside metabolic pathway, leading to the accumulation of downstream products, including nucleosides and purine bases.

### Autophagy limits obesity-induced xanthine/hypoxanthine secretion from adipocytes

Based on the intracellular changes in nucleotide metabolism, we next investigated whether this would have paracrine or endocrine effects on the tissue environment and systemically. Cumulative xanthine and hypoxanthine levels accumulate in mouse serum over the time course of HFD feeding (Fig 3A). Strikingly, serum xanthine and hypoxanthine were further elevated in the absence of adipocyte autophagy (Fig 3B), reflecting the intracellular metabolic rewiring in *Atg7^Ad^*adipocytes. Since we did not observe a shift of purine catabolism towards increased synthesis of its end product uric acid in *Atg7^Ad^* adipocytes, we postulated that xanthine might be rather than converted to uric acid intracellularly, actively secreted into the extracellular milieu by adipocytes. To understand whether these nucleobases were adipocyte-derived and controlled by autophagy, we performed targeted metabolomics on the adipocyte secretome. Similar to their intracellular levels, we found a notable accumulation of cytidine, uridine, guanosine, and xanthine in the secretome derived from *Atg7^Ad^* compared to WT adipocytes (Fig 3C). Xanthine and hypoxanthine levels more than doubled in the secretome of obese *Atg7^Ad^* adipocytes (Fig 3D), and we observed a strong negative correlation between the activity of the autophagy pathway and their secretion from obese gWAT in WT and *Atg7^Ad^* mice (Fig 3E). Purine nucleoside phosphorylase (PNP) plays a key role in purine catabolism, limiting the production of purine nucleobases (Korycka et al., 2007). Inhibition of PNP activity by the clinically approved drug forodesine resulted in a significantly lower xanthine and hypoxanthine secretion from both WT and *Atg7^Ad^*adipocytes, with the latter secretion being reduced to WT levels upon PNP inhibition (Fig 3F). Furthermore, inhibition of gWAT apoptosis by pan-caspase inhibitor Q-VD-OPh did not impact xanthine secretion from either WT nor *Atg7^Ad^* adipocytes (Fig 3G). Taken together, these interventional assays suggest that adipocytes actively generate xanthine and hypoxanthine through PNP activity and that their secretion is not a consequence of release from dying cells. These results altogether demonstrate that autophagy regulates xanthine and hypoxanthine secretion from visceral adipocyte during obesity.

**Figure 3:**
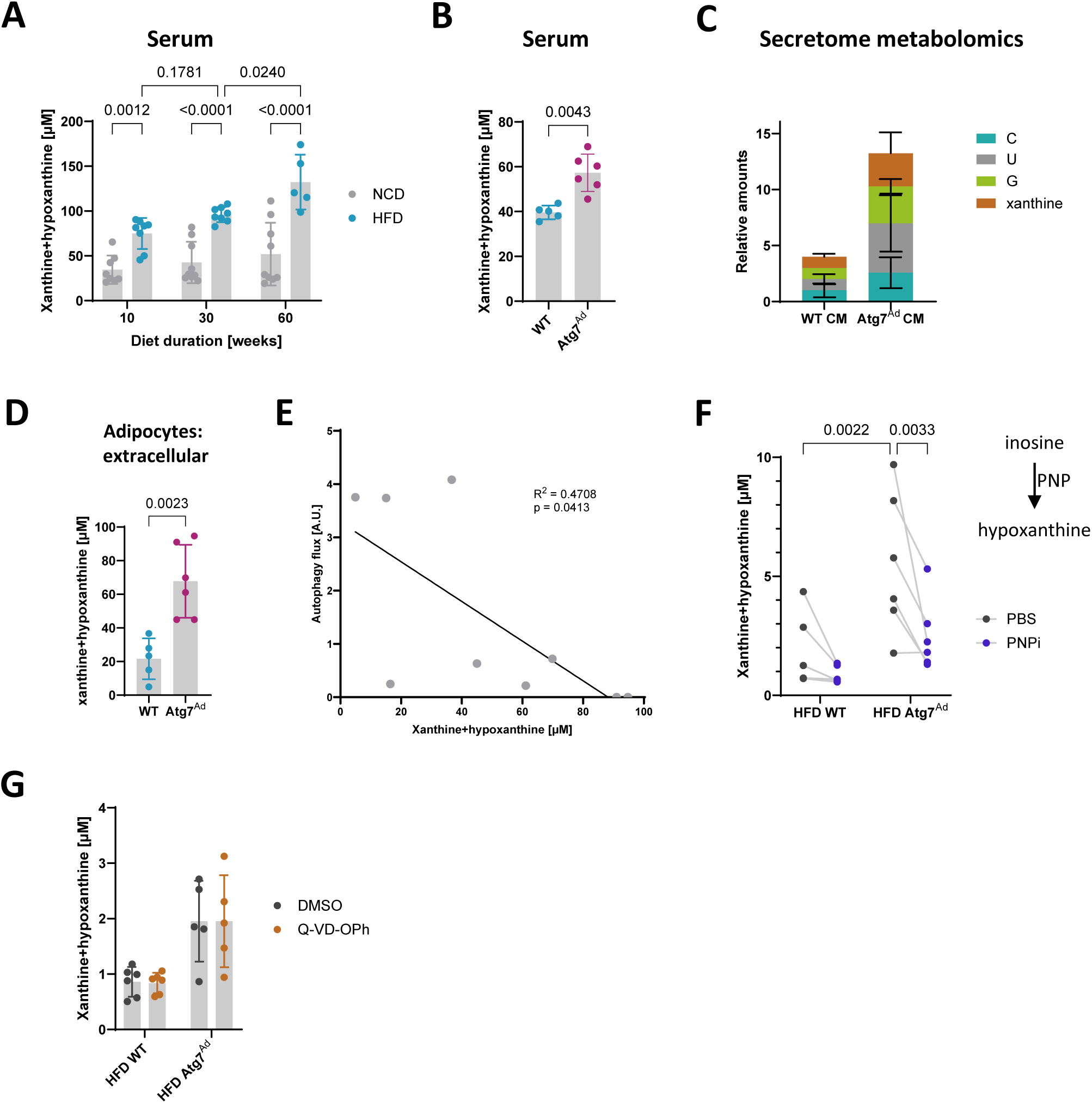
Elevated secretion of xanthine in response to obesity by adipocytes is limited by autophagy. A) Concentration of serum xanthine and hypoxanthine in WT mice were fed NCD or HFD for 10, 30 or 60 weeks. n = 5 – 8 mice. Data are merged from 3 independent experiments. B) Concentration of serum xanthine and hypoxanthine in WT and *Atg7^Ad^* mice fed with HFD for 16 weeks. n = 5-6 mice. Representative of 3 independent experiments. C) Relative abundance of nucleosides secreted by gWAT adipocytes isolated from WT and *Atg7^Ad^* mice following HFD feeding for 16 weeks and measured in metabolomics analysis. n = 2-3 mice. D) Concentration of xanthine and hypoxanthine secreted from gWAT adipocytes cultured over 24 hours *ex vivo*. Adipocytes were isolated from WT and *Atg7^Ad^* mice fed with HFD for 16 weeks. n = 5-6 mice. Representative of 3 independent experiments. E) Correlation analysis of the level of autophagy flux in gWAT and concentration of xanthine and hypoxanthine secreted from gWAT adipocytes as in (D). Adipocytes and gWAT were isolated from WT and *Atg7^Ad^* mice fed with HFD for 16 weeks. Western blot analysis of autophagy flux was calculated as (LC3-II (Inh) – LC3-II (Veh)). n = 9 mice. Data are merged from 3 independent experiments. F) Concentration of xanthine and hypoxanthine secreted from gWAT adipocytes cultured over 24 hours *ex vivo* and treated with either PBS or 10 µM forodesine, a purine nucleoside phosphorylase (PNP) inhibitor. Adipocytes were isolated from WT and *Atg7^Ad^* mice fed with HFD for 16 weeks. n = 6 mice. Representative of 3 independent experiments. G) Concentration of xanthine and hypoxanthine secreted from gWAT explants cultured overnight *ex vivo* and treated with either DMSO or 20 µM Q-VD-OPh, a pan-caspase inhibitor. gWAT was isolated from WT and *Atg7^Ad^*mice fed with HFD for 16 weeks. n = 5-6 mice. Representative of 3 independent experiments. Data are presented as mean ± SD. Dots represent individual biological replicates. Statistical analysis by two-way ANOVA with Tukey multi comparisons (A) or Fisher test (F), Mann-Whitney test (B), unpaired t-test (D) or Pearson R correlation analysis (E).

### Adipocyte autophagy controls WAT remodelling by limiting immune cell expansion

Given the dysregulated intra- and extracellular purine nucleoside signalling in *Atg7^Ad^* mice and the pronounced fibrosis, we set out to investigate whether these factors impact broader tissue remodelling and have a body-wide impact. We found that WAT body distribution exhibited remarkable differences with a reduced gWAT but expanded inguinal WAT (iWAT) deposition in obese *Atg7^Ad^* mice (Fig S4A). This was independent of weight gain or energy consumption between WT and *Atg7^Ad^* mice on either control (NCD) or high fat diet (HFD), where no differences were observed (Fig S4B-C). Deposition of fat into the visceral/gonadal area has been recognized to be more detrimental for obesity pathology and it has been suggested that iWAT expansion could potentially buffer the deleterious effects of visceral fat increase (Marcelin et al., 2022, González-Muniesa et al., 2017). In line with this hypothesis, changes in fat deposition in obese *Atg7^Ad^* mice were associated with improved obesity-induced metabolic syndrome, as demonstrated by increased glucose tolerance and lessened ectopic fat deposition in the liver (Fig S4D-G). Yet, no reduction in serum triglycerides and HDL cholesterol was observed (Fig S4H-I). In addition, obese *Atg7^Ad^* mice displayed no significant changes in circulating levels of adiponectin or leptin compared to controls (Fig S4J-K). Furthermore, autophagy-deficient adipocytes displayed no notable differences in cell size in gWAT (Fig S4L). These data suggest that autophagy-mediated adipocyte metabolic and tissue structural remodelling impact fat distribution in the pathological visceral WAT, alleviating obesity-induced metabolic syndrome.

Excessive ECM deposition in most tissues is commonly associated with increased secretion of pro-fibrotic cytokines that act to modulate the activity of ECM remodelling cells, such as transforming growth factor β (TGFβ) and osteopontin (OPN) (Gliniak et al., 2023, Meng et al., 2016, Icer and Gezmen-Karadag, 2018). To assess the production and release of these cytokines, we cultured gWAT *ex vivo* for six hours and measured their secretion. Both TGFβ and OPN secretion were increased in *Atg7^Ad^* obese mice (Fig 4A-B). The increase in these two cytokines was associated with a significant elevation in nuclear density in *Atg7^Ad^* gWAT (Fig 4C-D). This nearly 3-fold difference in nuclear density could not be attributed to adipocyte hyperplasia, as the number of unilocular adipocytes in the tissue remained constant (Fig 4E). The fibroblast population (PDGFRα^+^), the main cell type involved in ECM dynamics, decreased in the *Atg7^Ad^* gWAT (Fig 4G and Fig S5A). This observation was further supported by the expression of *Acta2*, a gene largely restricted to myofibroblasts, which was not increased in gWAT of obese *Atg7^Ad^* mice (Fig 1G). The endothelial cell population (CD31^+^) remained steady (Fig 4H and Fig S5A). Notably, however, the immune cell (CD45^+^) population, which can also be implicated in ECM dynamics (Marcelin et al., 2022), expanded significantly within the stromal vascular fraction of the *Atg7^Ad^* gWAT (Fig 4I and Fig S5A).

**Figure 4:**
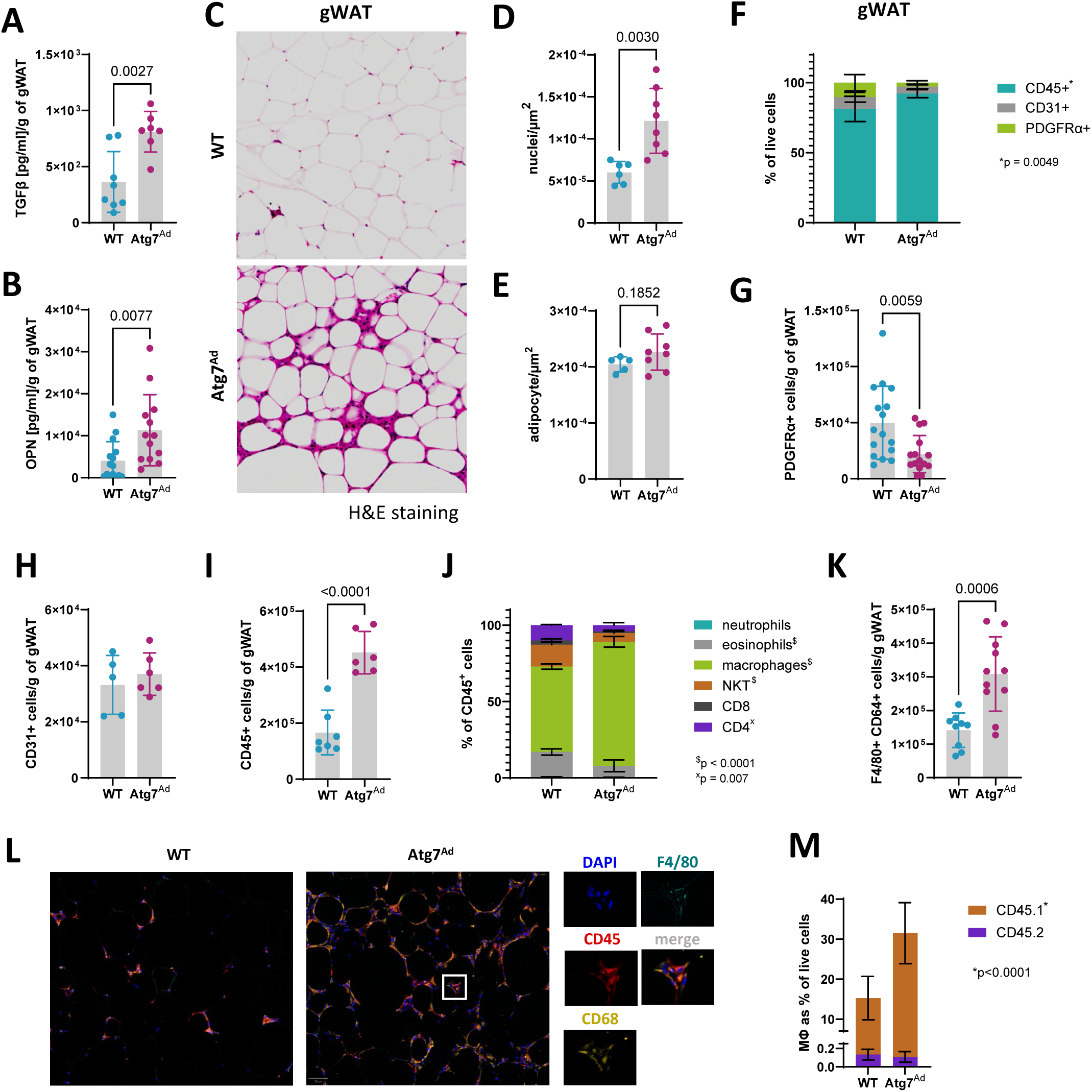
Loss of adipocyte autophagy results in macrophage infiltration. WT and *Atg7^Ad^* mice were fed HFD for 16 weeks before gWAT was isolated for analysis. A-B) Secretion of TGFβ (A) and osteopontin (OPN) (B) measured by ELISA. n = 7-15 mice. Data are merged from 3 independent experiments. C) H&E staining of gWAT. Scale bar, 200 µm. Representative of 3 independent experiments. D-E) Quantification of nuclei and adipocyte number from C. n = 5-8 mice. Representative of 3 independent experiments. F) Flow cytometry analysis of CD45^+^, CD31^+^, and PDGFRα^+^ populations in gWAT. n = 5 mice. Representative of 3 independent experiments. G-I) Absolute numbers of CD45^+^ (G), CD31^+^ (H), and PDGFRα^+^ (I) populations normalized to gram of WAT as in F. n = 5-16 mice. Data are merged from 3 independent experiments. J) Flow cytometry analysis of immune cell (CD45^+^) composition. NKT = natural killer T cell. n = 3-7 mice. Representative of 3 independent experiments. K) Flow cytometry analysis of F4/80^+^ CD64^+^ macrophage number in gWAT. n = 10 mice. Data are merged from 3 independent experiments. L) Representative immunofluorescence staining of F4/80, CD45 and CD68 of gWAT sections from WT and *Atg7^Ad^* mice following HFD feeding for 16 weeks. M) Flow cytometry analysis of F4/80^+^ CD64^+^ macrophage frequency labelled with CD45.1 (donor) and CD45.2 (host) congenic markers in gWAT after adoptive transfer. CD45.1 bone marrow cells were transferred in CD45.2 WT and *Atg7^Ad^*hosts, where conditional knockout was induced 25 days following the transfer and mice were fed HFD for an additional 12 weeks. n = 6 mice. Representative of 3 independent experiments. Data are presented as mean ± SD. Dots represent individual biological replicates. Statistical analysis by unpaired t-test (A, B, D, E, G, H, I, K) and two-way ANOVA with Šídák multi comparisons test (F, J, M).

Assessing the composition of the CD45^+^ compartment in gWAT by flow cytometry revealed that macrophages were the most prevalent population upon obesity, which further increased in abundance in *Atg7^Ad^* mice (Fig 4J and Fig S5B). The increase in macrophage numbers was confirmed by in *in-situ* imaging (Fig 4L), with macrophage numbers more than doubling (Fig 4K). To test whether these cells were derived from tissue-resident macrophages or infiltrating monocytes, we transplanted congenic bone marrow into WT or *Atg7^Ad^* hosts. Reconstitution of *Atg7^Ad^* mice with congenic CD45.1 bone marrow revealed that the majority of tissue-infiltrating macrophages were monocyte-derived and of those, there were twice as many in the obese adipose tissue of the *Atg7^Ad^*host compared to the WT host (Fig 4M). Collectively, these observations highlight the critical role of adipocyte autophagy in modulating the inflammatory environment during obesity by controlling macrophage infiltration.

### ATMs switch to a tissue-reparative phenotype in *Atg7^Ad^* gWAT

To better understand the identity and function of accumulated macrophages in *Atg7^Ad^* gWAT, we isolated F4/80^+^ CD64^+^ macrophages from gWAT of WT and *Atg7^Ad^* mice by fluorescence-activated cell sorting (FACS) and performed transcriptomics. Surprisingly, gene enrichment analysis of significantly dysregulated genes (Fig 5A) revealed that loss of adipocyte autophagy induces downregulation of pathways associated with inflammation and cytokine production (Fig 5B) while upregulating proliferative and tissue-remodelling processes as well as purine/nucleotide metabolism in ATMs (Fig 5C). To validate the results, we first cultured ATMs from gWAT of WT and *Atg7^Ad^* mice *ex vivo* and measured their secreted cytokines. We confirmed that macrophages from obese *Atg7^Ad^*mice notably decreased their cytokine production of IL-1β, IL-6, TNFα, and IL-10 (Fig 5D-G). In contrast, ATMs from obese *Atg7^Ad^*mice increased transcription of key pro-fibrotic tissue remodelling genes *Col3a1* and *Mmp14* (Fig 5H).

**Figure 5:**
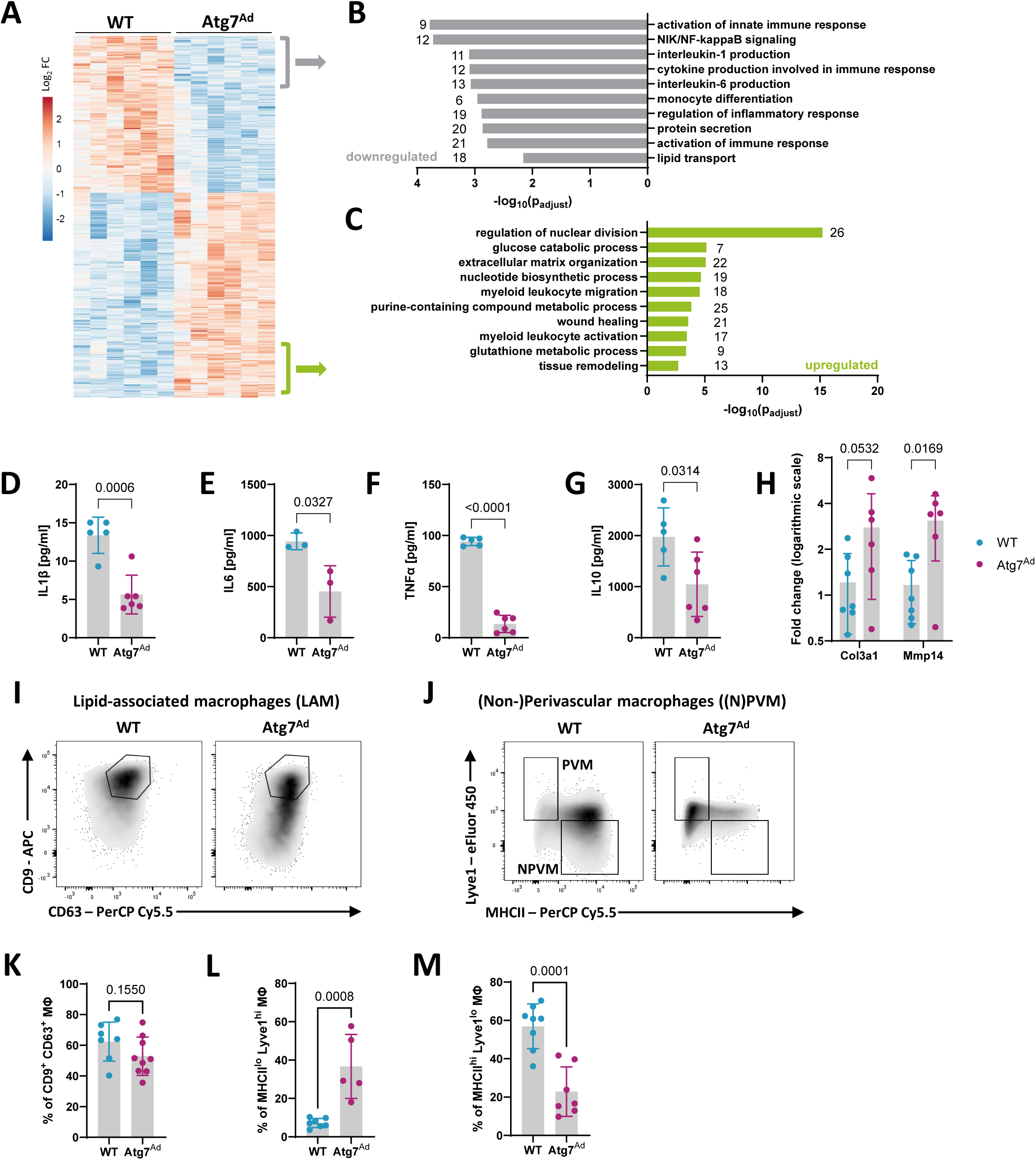
Macrophages acquire a tissue-reparative phenotype upon autophagy loss in gWAT adipocytes. A-C) Transcriptomics analysis of F4/80^+^ CD64^+^ macrophages isolated from gWAT of WT and *Atg7^Ad^* mice fed with HFD for 16 weeks. Hierarchical clustering of transcriptional profiles of the top 1000 differentially expressed genes (A). Colour coding represents the log2 fold difference between WT and *Atg7^Ad^* mice. Enrichment gene ontology (GO) analysis of downregulated (B) or upregulated (C) pathways in macrophages isolated from *Atg7^Ad^* compared to WT gWAT. The number of genes identified for each term is labelled. n = 6 mice. D-G) Secretion of IL-1β (D), IL-6 (E), TNFα (F), and IL-10 (G) by macrophages enriched from gWAT of WT and *Atg7^Ad^*mice fed HFD for 16 weeks. n = 3-6 mice. Representative of 3 independent experiments. H) Relative mRNA levels of extracellular matrix (ECM)-related genes in sorted F4/80^+^ CD64^+^ macrophages isolated from gWAT of WT and *Atg7^Ad^* mice fed HFD for 16 weeks measured by qRT-PCR. Data presented as log2 fold difference. n = 6-7 mice. Representative of 3 independent experiments. I-M) Representative plots of lipid-associated macrophages (LAM) (I), identified as CD63^+^ CD9^+^, perivascular (PVM) and non-perivascular macrophages (NPVM) (J), identified as MHCII^low^ Lyve1^high^ and MHCII^high^ Lyve1^low^, respectively, assessed by flow cytometry. Quantification of LAM (K), PVM (L) and NPVM (M) frequency in the gates shown. n = 5-9 mice. Data are merged from 3 independent experiments. Data are presented as mean ± SD. Dots represent individual biological replicates. Statistical analysis by unpaired t-test (D-G, K-M) and two-way ANOVA with Šídák multi comparisons test (H).

The growing recognition of ATM plasticity and heterogeneity has revealed a complexity that renders the traditional M1/M2 (pro- and anti-inflammatory) paradigm overly simplistic and outdated (Maniyadath et al., 2023, Nance et al., 2022). The recent classification obtained under both normal chow and high fat diet using single-cell RNA sequencing (scRNA-Seq) suggests three main macrophage subtypes, including perivascular-like macrophages (PVM), non-perivascular-like macrophages (NPVM), and lipid-associated macrophages (LAM) (Hill et al., 2018, Jaitin et al., 2019, Chakarov et al., 2019, Xu et al., 2013, Sarvari et al., 2021). While NPVMs and LAMs mediate inflammatory processes, PVMs control tissue repair (Matz et al., 2023). Analysis of these macrophage populations by flow cytometry (Fig 5I-J) revealed no difference in LAM (marked as F4/80^+^ CD64^+^ CD9^+^ CD63^+^) abundance between obese *Atg7^Ad^* and WT gWAT (Fig 5I, K). In contrast, we found tissue reparative PVM (marked as F4/80^+^ CD64^+^ Lyve1^high^ MHCII^low^) more than seven-fold increased and antigen-presenting NPVM (marked as F4/80^+^ CD64^+^ Lyve1^low^ MHCII^high^) three-fold decreased among macrophages isolated from *Atg7^Ad^* gWAT (Fig 5J, L-M). In summary, we uncovered that in gWAT of obese *Atg7^Ad^* mice, macrophages switch from a predominantly pro-inflammatory to a tissue-reparative pro-fibrotic phenotype, which is accompanied by a strong ECM transcriptional signature.

### Metabolic dysregulation of *Atg7^Ad^* adipocytes is signalled through xanthine and hypoxanthine to macrophages for a tissue-reparative phenotypic switch

Observing that autophagy significantly impacted adipocyte purine nucleoside metabolism, which might, in turn, influence the surrounding microenvironment, we aimed to determine whether purine nucleosides could induce a tissue reparative phenotype in macrophages. In pursuit of this goal, we first tested whether the adipocyte secretome could switch macrophages *in vivo* by cultivating ATMs isolated from lean adipose tissue in the presence of the secretome derived from either obese WT or *Atg7^Ad^* adipocytes. Three days after the exposure, we observed a significant increase in the Lyve1^high^ MHCII^low^ tissue repair macrophage population as well as the upregulation of ECM-related genes *Col3a1*, *Mmp14*, and *Timp1* in macrophages exposed to *Atg7^Ad^* adipocyte-derived secretome (Fig 6A-C). These results mimicked our observations *in vivo*, suggesting that adipocyte-derived soluble signals are responsible for the macrophage phenotype.

**Figure 6:**
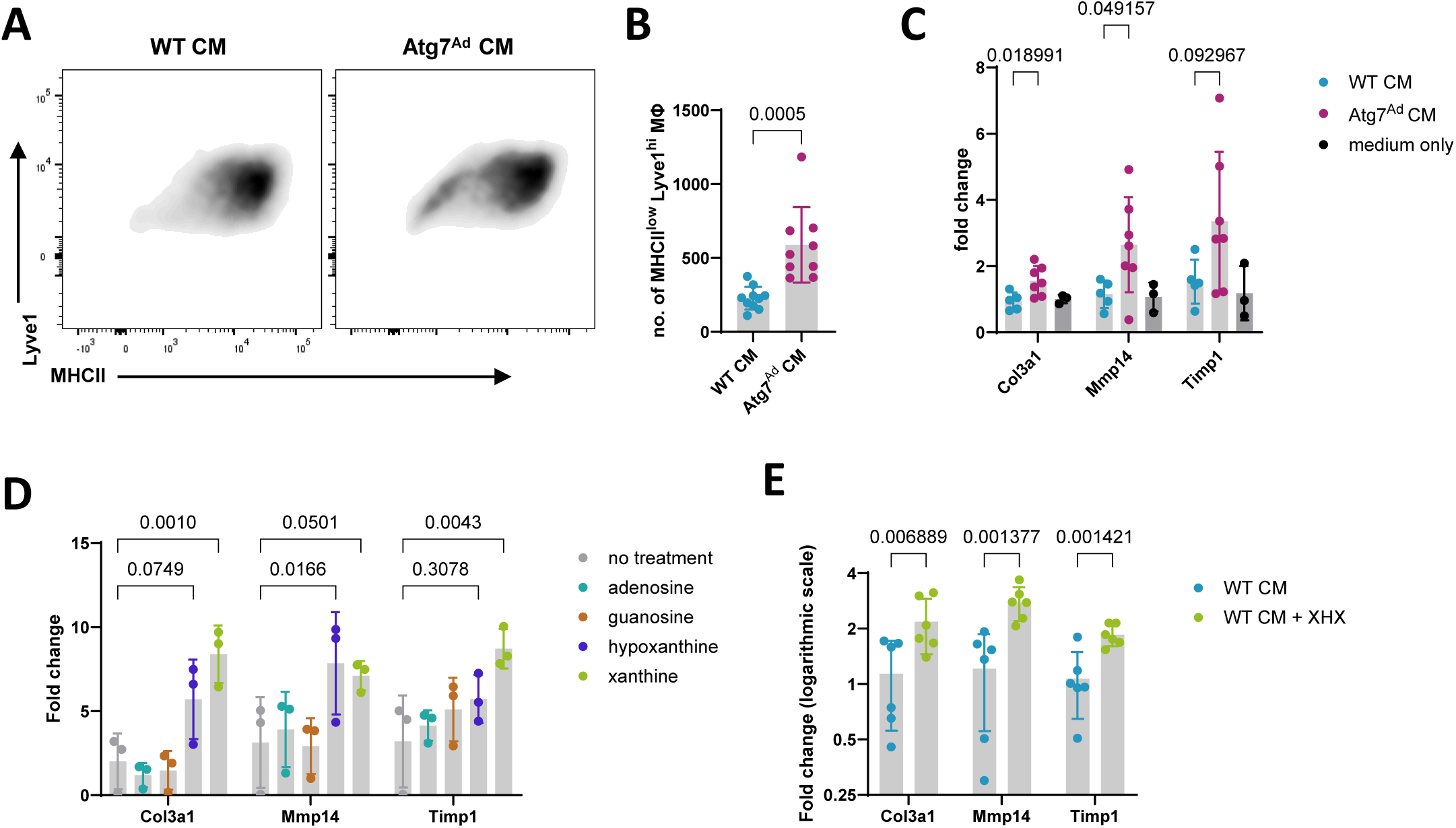
Autophagy in obese adipocytes inhibits tissue-reparative macrophages and fibrosis via purine nucleoside signalling. A-C) Macrophages were isolated from lean WT gWAT and cultivated *in vitro* in the presence of conditioned medium (CM) generated by 24-hour *ex vivo* incubation of obese adipocytes macrophages assessed by flow cytometry (A). Quantification of flow cytometry analysis of MHCII^low^ Lyve1^high^ F4/80^+^ CD64^+^ macrophage number after 72 hours of treatment with CM from WT or *Atg7^Ad^* adipocytes (B). Relative mRNA levels of ECM-related genes in macrophages after 72 hours of treatment with CM from WT or *Atg7^Ad^* adipocytes or baseline full medium (C). RNA levels measured by qRT-PCR. n = 3-10 mice. Representative of 3 independent experiments. D) Macrophages were isolated from lean WT gWAT and cultivated *in vitro* for 72 hours in baseline full medium supplemented with 50 ng/ml of M-CSF and 100 µM of either adenosine, guanosine, hypoxanthine or xanthine. Relative mRNA levels of ECM-related genes were measured by qRT-PCR. n = 3 mice. Representative of 3 independent experiments. E) Relative mRNA levels of ECM-related genes in macrophages after 72 hours of treatment with CM from obese WT adipocytes with or without 100 µM supplementation of both xanthine and hypoxanthine (XHX). The macrophages were isolated as in (A-C) and RNA levels were measured by qRT-PCR. Data presented as log2 fold difference. n = 6 mice. Representative of 3 independent experiments. Data are presented as mean ± SD. Dots represent individual biological replicates. Statistical analysis by unpaired t-test (A), multiple unpaired t-tests (B, D), or two-way ANOVA with Dunnett’s multiple comparisons test (C).

We next aimed to determine whether purine nucleosides could be responsible for these observations. To this end, lean ATMs were cultured *in vitro* for 72 hours in 50 ng/ml of M-CSF supplemented with 100 µM of either adenosine, guanosine, hypoxanthine or xanthine (Fig 6D). Xanthine, and to a lesser extent hypoxanthine, led to a significant upregulation of ECM-related genes, whereas adenosine and guanosine did not. To further test whether hypoxanthine and xanthine could indeed trigger a tissue-reparative switch, lean ATMs were treated *in vitro* with the secretome of obese WT adipocytes for 72 hours supplemented with a mixture of 100 µM xanthine and hypoxanthine each. We observed a marked increase in the pro-fibrotic signature genes *Mmp14*, *Col3a1*, and *Timp1* (Fig 6E). Collectively these results suggest that increased release of xanthine and hypoxanthine can promote tissue-reparative switch in macrophages during obesity. Thus, adipocyte autophagy presents a critical point during obesity that controls tissue inflammation versus repair balance via control of purine nucleoside metabolism and its extracellular signals.

## DISCUSSION

In this study, we have identified autophagy as a major brake on WAT fibrosis. Combining genetic model and dietary intervention with proteomic, metabolomic, and functional analyses we uncovered a critical role of autophagy in supporting adipocyte metabolic needs during excessive growth, limiting purine nucleoside catabolism. By studying the nucleoside metabolic changes upon loss of autophagy, we identified (hypo)xanthine-driven adipocyte-to-macrophage crosstalk. Finally, our work revealed a critical role of autophagy in limiting WAT ECM pathological remodelling through (hypo)xanthine-induced macrophage tissue repair phenotype.

Understanding the changes in autophagy activity in adipose tissues during obesity in both humans and mice remained elusive, despite numerous reports (Clemente-Postigo et al., 2020, Jansen et al., 2012, Kosacka et al., 2015, Kovsan et al., 2011, Mizunoe et al., 2017, Nuñez et al., 2013, Öst et al., 2010, Soussi et al., 2016, Soussi et al., 2015). In addition, the lack of clarity on the mechanism and function of autophagy in obese WAT highlighted the complex and poorly understood role of autophagy. While it has been reported that adipocyte autophagy supports adipose tissue-liver crosstalk, contradictory conclusions were drawn in the different studies (Cai et al., 2018, Sakane et al., 2021). Our data suggest that obesity dysregulates autophagy both in humans and mice and that autophagy primarily increases with obesity in mice, with an eventual drop after prolonged high fat diet feeding, perhaps explaining a few studies that showed decreased autophagy levels with obesity (Soussi et al., 2016, Soussi et al., 2015). We find that the primary function of autophagy is the support of high metabolic demands of adipocytes during fat mass expansion. Adipocyte metabolism underlying fat storage and turnover is well understood (Morigny et al., 2021), and is majorly determined by an increase in WAT mass. Nevertheless, our understanding of metabolic rewiring beyond glucose and lipid metabolism remains limited, with only scarce evidence for the role of other key metabolic processes in adipocytes (Nagao et al., 2017, Park et al., 2017, Kather, 1990). We find that in humans, purine nucleoside metabolism represents one of the main dysregulated metabolic pathways in obese adipocytes. Our proteomics and metabolomics analyses revealed that autophagy critically supports nucleotide and amino acid pools in obese adipocytes. Similar autophagy-dependent changes have been previously observed in lung cancer cells under starvation and haematopoietic stem cells (Guo et al., 2016, Borsa et al., 2024, Zhang et al., 2018). Similar to these cell types, mature adipocytes have a highly dynamic metabolic demand and can enter a pseudo-starvation state through adipokine signalling in obesity (James et al., 2021). Furthermore, increased production of purine nucleoside catabolic intermediates, such as hypoxanthine and xanthine, has been previously suggested to relate to ATP depletion (Guo et al., 2016, Harkness, 1988, Harkness et al., 1983, Harkness and Lund, 1983), which we also observed. Therefore, we believe these observations spanning several different cell types share a common molecular mechanism and highlight the indispensable role of autophagy in the provision of bioenergetic and biosynthetic substrates, responding to stress, maintaining redox homeostasis and survival.

Failure of adipocyte autophagy induction resulted in gWAT fibrosis, which is the more fibrosis-prone WAT depot (Marcelin et al., 2022). The role of autophagy in fibrosis is controversial and highly context-dependent (Li et al., 2020, Sun et al., 2021). While the relationship between autophagy and adipose tissue fibrosis has not been experimentally addressed to date, their potential link has been proposed recently (Oh et al., 2023). Tissue fibrosis develops when either ECM deposition or turnover become dysfunctional and is difficult to reverse (Reggio et al., 2013). Fibrosis of adipose tissue has been traditionally seen as detrimental as it mechanically stiffens the tissue, thereby negatively impacting its critical plasticity feature in the response to nutrient status (Gliniak et al., 2023). Nevertheless, fibrosis is an essential component of tissue repair that limits tissue damage and aims to restore functional tissue architecture, improve recovery, and survival (Henderson et al., 2020). In a chronic setting, when damage is persistent or severe, however, fibrosis leads to disruption of tissue architecture, interferes with organ function, and can ultimately lead to organ failure (Medzhitov, 2021). We observed a chronic increase in fibrosis of gWAT over the time of HFD feeding in

normal adipose tissue but much accelerated in adipose tissue without autophagy. We suggest that initially, fibrosis acts to prevent acute and excessive tissue damage due to impaired adipocyte homeostasis and function upon autophagy depletion. Eventually, however, chronic accumulation of ECM likely leads to a broader adipose tissue dysfunction. Similar observations have recently been made in the pancreas (Baer et al., 2023). Since WAT is not functionally compartmentalized, the detrimental effects of chronic fibrosis at the organismal level are difficult to discern. Indeed, increased deposition in the subcutaneous area positively correlates with more favourable disease outcomes compared to visceral deposition (Sakers et al., 2022). It has been proposed that the expansion of subcutaneous WAT could potentially help reduce the detrimental impact of visceral WAT expansion (Marcelin et al., 2022). Concomitant with this, we observed the physical limitation of the pathological visceral WAT expansion by fibrosis, improving glucose homeostasis, and reducing ectopic fat deposition to the liver. Understanding the determinants of WAT remodelling and fibrosis holds an important therapeutic potential to improve obesity management and health outcomes of obese patients.

Excessive pericellular fibrosis positively correlated with a pronounced accumulation of tissue-reparative macrophages. Increased macrophage accumulation in adipocyte autophagy-deficient gWAT has been observed before but never studied in detail (Cai et al., 2018, Sakane et al., 2021). Macrophages are known as key regulators of tissue repair, regeneration, and fibrosis (Lech and Anders, 2013), and this may be true for adipose tissues as well (Marcelin et al., 2022, Vila et al., 2014). The evidence, however, remains scarce, with elastin and TLR4 signalling being proposed to play a role in macrophage-induced WAT fibrosis during obesity (Martinez-Santibanez et al., 2015, Vila et al., 2014). On the other hand, ATMs have also been proposed to prevent pathological changes of ECM and limit the development of gWAT fibrosis (Chen et al., 2021). Nevertheless, it remains unclear which signals induce the macrophage pro- or anti-fibrotic phenotypic switch that could serve as important balance checkpoints and therapy targets in fibrotic diseases. Local metabolic signals contributing to immune cell fates are becoming an area of increasing interest (Bacigalupa et al., 2024, Richter et al., 2018). We show here for the first time that products of nucleoside catabolism, xanthine and hypoxanthine, can act as determinants of adipose tissue macrophage fate, resulting in a tissue-reparative phenotype. Notably, we found xanthine and hypoxanthine increased with obesity progression in mouse serum, and similar observations have been made before in human, identifying adipose tissue as one of the main contributing factors (Nagao et al., 2018, Furuhashi et al., 2020, Ho et al., 2016, Xie et al., 2014). Furthermore, adipocytes have been previously described to actively release nucleosides upon stress, including hypoxanthine, xanthine, inosine, guanosine, and uridine (Deng et al., 2018, Pfeifer et al., 2024, Fromme et al., 2018, Kather, 1988, Kather, 1990). We uncover for the first time that autophagy acts as a brake for the active release of nucleosides and nucleobases. A directed secretion of xanthine by T cells has been identified to relay cell-extrinsic effects under stress conditions (Fan et al., 2019). These results together with our observations suggest a common molecular signalling mechanism of cellular stress to the microenvironment via purine nucleobase signals. Our data indicate that this extracellular purine signal can be controlled by autophagy, which helps the cell to adapt to novel metabolic challenges such as excessive storage of fat. It is plausible that by activating nucleotide degradation, autophagy-deficient adipocytes salvage NADPH or ribose through the PPP. This enables them to partially sustain their metabolism and oxidative stress, sourcing carbon for energy, antioxidant molecules, and anabolic precursor generation. In turn, nucleotide catabolites signal the altered adipocyte state to the macrophages, which by remodelling ECM shut down the tissue and limit systemic dysregulation. While autophagy-dependent metabolic signals have been previously reported to play a role in cancer and inflammatory bowel disease (Sousa et al., 2016, Poillet-Perez et al., 2018, Richter et al., 2023), pro-fibrotic purine nucleoside catabolites have not been described before.

In conclusion, our work highlights the key role of autophagy acting as a brake in the control of adipocyte nucleoside metabolism and tissue integrity in diet-induced obesity. When dysfunctional, this leads to uncontrolled activation of metabolic rewiring that generates purine catabolites xanthine and hypoxanthine, signalling tissue repair. By depleting autophagy, we uncover a purine nucleobase-mediated pro-fibrotic signalling pathway, and further research is necessary to elucidate whether these signalling molecules control fibrosis of other tissues and organs, potentially deeming them druggable targets.

## ACKNOWLEDGEMENTS

We thank Patricia Cotta Moreira, Luke Barker, Emily Wyeth, Daniel Andrew, and Mino Medghalchi from the Biomedical Services for their responsible care and assistance with animal well-being. Histology was performed with the help of the Kennedy Institute Histology Facility, with special thanks to Dr. Ida Parisi. Dr Johanna ten Hoeve-Scott at UCLA Metabolomics Center, Dr Ulrike Brüning and Dr Jennifer Kirwan at BIH Charité Berlin for their help with metabolomics sample analysis. Dr. Moustafa Attar for his help with the experimental design of the transcriptomics experiment. Jonathan Webber for his help with the flow cytometry experimental design. Cell DIVE Facility team at the Kennedy Institute of Rheumatology for their help with immunofluorescent staining. This work was supported by grants from the Wellcome Trust to A.K.S. (Investigator award 220784/Z/20/Z) and M.B. (Sir Henry Wellcome Fellowship 220452/Z/20/Z), the Kennedy Trust for Rheumatology Research (KTTR) Studentship to K.P. (KEN192001) and J.K.L.K. (awarded to Marco Fritzsche), The Kenneth Rainin Foundation to A.K.S. (20220003/20230038), Clarendon Fund Scholarship to K.P., Medical Research Council Doctoral Training Partnership Grant (BRT00030) to K.P., Ramage Scholarship to K.P., PhD studentship award 203803/Z16/Z to F.C.R., Versus Arthritis grant 22617 to G.A ., EPA Cephalosporin Fund to J.K.L.K., EMBO Postdoctoral Fellowship (ALTF115-2019) to A.V.L.V. Li-cor Odyssey imager was funded by ERC AdG 670930. Flow cytometry and microscopy facilities were supported by KTTR. Graphical summaries were created with BioRender.com.

## AUTHOR CONTRIBUTIONS

Conceptualization: K.P., F.P., G.A., and A.K.S. Methodology, investigation, analysis, visualization, and validation: K.P., A.H.K. F.C.R., M.B., A.V.L-V., O.P., S.G., K.K., K.Z., M.K., L.K., and G.A. Essential reagents and support: O.P., L.K., P.M. and S.S. Writing of original draft: K.P., G.A., and A.K.S. Funding acquisition, supervision, and project administration: F.P., G.A., and A.K.S. Editing of draft: all authors.

## METHODS

### Lead contact

Information and requests for reagents and resources should be directed to the Lead Contact, Anna Katharina Simon.

### Mouse models

Adipoq-Cre^ERT2^ mice (Sassmann et al., 2010) were purchased from Charles River, UK (JAX stock number: 025124) and crossed to Atg7^fl/fl^ mice (Komatsu et al., 2005). Genetic deletion was induced at 6-8 weeks of age by oral gavage of 4 mg tamoxifen per mouse for five consecutive days. Tamoxifen was given to all groups of mice. Two days after receiving the last tamoxifen dose, mice were subjected to an altered diet regime with either a high fat diet with 60 kcal% fat (D12492i, Research Diets) or a complementary normal chow diet with 10 kcal% fat (D12450Ji, Research Diets) for the duration stated in the text. Wild-type C57BL/6J or B6.SJL.CD45.1 mice were bred in-house. Experimental cages were sex- and age-matched and balanced for genotypes. All data shown except proteomics and metabolomics data are pooled from both sexes. Mice were maintained on a 12 h dark/light cycle and housed in groups of 3-5 with unlimited access to water and food under specific pathogen-free conditions. The temperature was kept between 20 and 24 °C, with a humidity level of 45–65%. All experiments were performed in accordance with approved procedures by the Local Review Committee and the Home Office under the project license (PPL30/3388 and P01275425).

### Bone marrow chimera generation

Recipient WT and Atg7^Ad^ mice were lethally irradiated with 11 Gray dose before intravenously injecting between 250-300,000 B6.SJL.CD45.1 donor cells (equal numbers in the same experiment to allow comparison between the two groups). Cell replenishment was followed bi-weekly. Five weeks after irradiation, mice were treated with tamoxifen for genetic deletion and fed high fat diet for a total of 12 weeks (as described above).

### Tissue processing, macrophage isolation and primary cell culture

Adipose tissue digestion was performed as previously described (Richter et al., 2023). In brief, depots were digested in DMEM containing 1 % fatty acid-free BSA, 5 % HEPES, 0.2 mg/ml Liberase TL (Roche), and 20 μg/mL DNaseI. Tissues were minced and incubated for 25-30 min at 37°C at 180 rpm. Digested tissue was strained through a 300 μm mesh and the digestion was quenched by the addition of PBS with 0.5 % BSA and 2 mM EDTA. Adipocyte and stromal vascular fraction were separated by 7 min centrifugation at 500 g and collected for further analysis.

To generate a conditioned medium, adipocytes were collected with wide-bore tips and washed three times with PBS. The floating fraction was collected and 250 µl of packed adipocytes were seeded in 500 µl of RPMI containing 10 % foetal bovine serum (FBS) and 1 % penicillin/streptomycin (P/S) and incubated for 24 hours at 37°C. Alternatively, the medium was supplemented with 10 µM forodesine. After incubation, the medium and cells were harvested, centrifuged at 300 g for 5 min and purified medium was collected and snap-frozen for *in vitro* experiments.

For primary cell culture, the stromal vascular fraction was enriched for CD11b^+^ cells with CD11b MicroBeads (Miltenyi Biotec) according to manufacturers’ instructions after red blood cell lysis. 350,000 cells were seeded in 24-well plates in RPMI containing 10 % FBS and 1 % P/S and incubated overnight to allow macrophages to attach. The following day, macrophages were enriched by washing the wells with room temperature PBS and treated with experimental conditions. For conditioned medium treatment, RPMI containing 10 % FBS and 1 % P/S was mixed with the conditioned medium in a 2:1 ratio and applied to the cells for 72 hours. For nucleoside treatment, macrophages were cultured in RPMI containing 10 % FBS and 1 % P/S and 50 ng/ml M-CSF and treated with 100 µM of either adenosine, guanosine, hypoxanthine or xanthine.

### Glucose tolerance test

Mice were subjected to 12 hours fast before measuring fasted glucose levels. To monitor response to bolus glucose, 1.5 g of glucose per kg of body mass was injected intraperitoneally. Blood glucose levels were measured via tail clip at 15, 30, 60, 90, 120, and 180 min after injection with a glucose meter (Freestyle Lite, Abbott).

### Histology and immunohistochemistry

Adipose tissues and liver were fixed in 10 % neutral buffered formalin for 24 and 48 hours, respectively. All tissues were transferred to 70 % ethanol and sent to the Kennedy Institute of Rheumatology Histology service for paraffin embedding, sectioning (5 µm), and staining. Haematoxylin and eosin (H&E) and picrosirius red staining were performed according to standard protocols. Images were acquired with a Zeiss Axioscan 7 scanning microscope. Image analysis and quantification were performed using QuPath, Image J, and an in-house developed script (available at https://github.com/Oxford-Zeiss-Centre-of-Excellence/pyHisto). In brief, the blind colour deconvolution method was used based on a stain vector estimation (Ruifrok and Johnston, 2001), followed by Otsu thresholding and determination of collage-to-area ratio. For immunofluorescence staining, WAT was fixed in 4 % paraformaldehyde for 24 hours, embedded and sectioned as above. Tissues were subsequently deparaffinized and heat-retrieved at 100°C for 20 min in Citrate antigen retrieval solution (Vector Laboratories, pH 6.0), and allowed to cool naturally in Tris antigen retrieval solution (10 mM, pH 8.8-9.0). Slides were washed in PBS and blocked overnight in 10 % donkey serum and 3 % BSA. The next day, slides were incubated with DAPI for 15 min to record a background scan. After background imaging, slides were incubated for 20 min with Fc block reagent (1:200 in 3 % BSA), followed by primary antibody incubation overnight at 4°C. Slides were washed in PBST, and incubated with secondary antibodies diluted in 3 % BSA for one hour at room temperature. Following washes in PBST, slides were mounted, and images were acquired with a GE Cell DIVE multiplex imager. See the Key resources table for a list of primary and secondary antibodies.

### Western blot

Autophagy flux in WAT was assessed by incubating adipose tissue in full DMEM supplemented with 100 nM Bafilomycin A1 and 20 mM NH4Cl for 4 h. DMSO was used as a ‘vehicle’ control. To determine apoptosis, WAT explants were incubated in full RPMI supplemented with either DMSO (vehicle control) or 20 µM Q-VD-OPh overnight. Protein extraction was performed as published (An and Scherer, 2020). Briefly, 500 μL of lysis buffer containing protease inhibitors and PhosphoStop was added per 100mg of tissue. Cells were lysed using Qiagen TissueLyser II and lipid contamination was removed through serial centrifugation. Protein content was determined using a BCA Protein Assay kit and 15 μg of protein was separated on a 4-12% Bis-Tris SDS PAGE gel. After wet-transferred onto a PVDF membrane, membranes were blocked in 5 % milk in TBST, and incubated with primary antibodies overnight. Proteins were visualized on a membrane using IRDye 800 or IRDye 680 (LI-COR Biosciences) secondary antibodies at the dilution 1:10,000 (LI-COR). Quantification was performed with ImageStudio software (LI-COR). Autophagic flux was calculated as: (LC3-II (Inh) – LC3-II (Veh)). See the Key resources table for a list of primary and secondary antibodies.

### Enzyme-linked immunosorbent assay (ELISA)

Adipose tissue secretome was generated by incubating 200 mg of adipose tissue explants (each explant ∼ 10 mg) in 1 ml of RPMI containing 10 % FBS and 1 % P/S for 6 hours at 37°C. Cytokine levels in supernatant were measured by commercially available ELISA kits. See the Key resources table for a list of ELISA kits. All cytokine levels were normalized to input tissue weight.

### Serum chemistry

After eight hours of fasting, serum was collected by a cardiac puncture, collected in Microtainer tubes, and centrifuged for 90 s at 15,000 g. Triglycerides and high-density lipoprotein were measured using a Beckman Coulter AU680 clinical chemistry analyser.

### Gene expression analysis (qRT-PCR)

Tissues were homogenized in TRI reagent (Sigma) with ceramic beads (Bertin Instruments) using a Precellys 24 homogenizer (Bertin Instruments). RNA was extracted using RNeasy Mini or Micro Kit (Qiagen). RNA yield and quality were assessed using a NanoDrop and cDNA was synthesized using a High-Capacity RNA-to-cDNA™ kit (ThermoFisher). Gene expression was measured using TaqMan Fast Advanced Master Mix on a ViiA7 real-time PCR system. Values were normalized to *Ppia* reference gene using the comparative Ct method. Primers are listed in the Key resources table.

### Flow cytometry and cell sorting (FACS)

Cells for flow cytometry staining were isolated as described above. For surface staining, cells were incubated with fluorochrome-conjugated antibodies, LIVE/DEAD Fixable Stains and Fc receptor block antibody for 20 min at 4°C. This was followed by a 10 min fixation with 4% PFA at room temperature. Samples were acquired on the Fortessa X-20 flow cytometer (BD Biosciences). Data were analysed with FlowJo v10.8.0. See the Key resources table for a list of flow cytometry antibodies.

### Transcriptomics (bulk RNA sequencing)

Macrophages were isolated by FACS and RNA was extracted as described above. To generate PolyA libraries, cDNA was end-repaired, A-tailed and adapter-ligated. Libraries were then size-selected, multiplexed, quality-controlled, and sequenced using a NovaSeq6000. Quality control of raw reads was performed with a pipeline readqc.py (https://github.com/cgat-developers/cgat-flow). The resulting reads were aligned to the GRCm38/Mm10 reference genome using the pseudoalignment method Salmon (Patro et al., 2017). DEseq2 (v1.38.3) was used for differential gene expression analysis (Love et al., 2014). The workflow included the estimation of size factors, dispersion estimation, and fitting of a negative binomial generalized linear model. Prior to differential expression analysis, batch effects attributed to sex were corrected using the limma package (v3.54.1) (Ritchie et al., 2015). Genes were considered significantly differentially expressed based on an adjusted p-value threshold of <0.05, after correcting for multiple testing using the Benjamini-Hochberg procedure. To explore the biological implications of the differentially expressed genes, gene set enrichment analysis was performed using the clusterProfiler package (v4.6.0). The transcriptomics dataset is available at GEO: GSE263837.

### Single nucleus RNA-seq analysis

The adipocyte dataset was downloaded from the GEO database (GSE176171), originally published by (Emont et al., 2022). The dataset was categorized into lean or obese groups based on BMI, following the methodology outlined in the source paper (Emont et al., 2022). To facilitate visualization, UMAP (Uniform Manifold Approximation and Projection) was recalculated, and further data analysis was conducted using the Seurat package for single-cell RNA-seq analysis. Functional enrichment analysis was performed using the ClusterProfiler package (v4.6.0).

### Proteomics

Adipocytes were isolated as a floating fraction upon digestion and lysed and digested in SDC buffer. Specifically, a pellet of 100 µl packed adipocytes was lysed in SDC-buffer containing 2% (w/v) sodium deoxycholate (SDC; Sigma-Aldrich), 20 mM dithiothreitol (Sigma-Aldrich), 80 mM chloroacetamide (Sigma-Aldrich), and 200 mM Tris-HCl (pH 8). After being heated at 95°C for 10 minutes, the lysates were digested enzymatically using endopeptidase LysC (Wako) and sequence grade trypsin (Promega) at a protein:enzyme ratio of 50:1. The digestion process occurred overnight at 37°C. For reversed-phase liquid chromatography coupled to mass spectrometry (LC-MS) analysis, each sample replicate was injected with 1 µg of peptide amount into an EASY-nLC 1200 system (Thermo Fisher Scientific) for separation, using a 110-minute gradient. Mass spectrometric measurements were carried out using an Exploris 480 (Thermo Fisher Scientific) instrument in data-independent acquisition (DIA) mode, which utilised an isolation scheme with asymmetric isolation window sizes. The raw files were analysed using DIA-NN version 1.8.1 (Demichev et al., 2020) in library-free mode, with a false discovery rate (FDR) cutoff of 0.01 and relaxed protein inference criteria, while employing the match-between runs option. The spectra were compared to a Uniprot mouse database (2022-03), which included isoforms. The protein intensities were normalised using MaxLFQ and filtered to ensure that each protein had at least 50% valid values across all experiments, with an additional filter to retain at least 3 valid values in at least one experimental group. The limma package (Ritchie et al., 2015) was used to calculate two-sample moderated t-statistics for significance calling. The nominal P-values were adjusted using the Benjamini-Hochberg method.

### Mass spectrometry

Adipocytes and conditioned medium were obtained as described above. Metabolite extractions from frozen cell pellets were performed through a two-phase extraction with 80 % MeOH and chloroform. In brief, when just thawed, cells were resuspended in 500 µl ice-cold 80 % MeOH, vortexed and sonicated in an ice bath for 6 min. Following one hour of incubation on dry ice, tubes were centrifuged at 16,000 g for 10 min at 4°C, and supernatants were mixed with water and chloroform in 1:1:1 ratio. Each sample was vortexed for 1 min and centrifuged at top speed for 15 min at 4°C. Finally, 600 μl of the top aqueous layer was transferred to a glass vial and evaporated using an EZ-2Elite evaporator (Genevac). Samples were stored at −80 °C before analysis. The BCA assay was performed on the airdried pellets, resuspended in 200 µl of 0.2 M NaOH and heated at 95°C for 20 min. Metabolite extractions from frozen conditioned medium precleared from cells and cell debris were performed after removing cells and cell debris by mixing 20 µl of clarified medium with 500 µl ice-cold 80 % MeOH/20 % H_2_O. After vortex and 30 min incubation at -80°C, samples were centrifuged at 16,000 g for 10 min at 4°C, transferred to a glass vial and evaporated using an EZ-2Elite evaporator (Genevac). Dried extracts were stored at −80 °C before analysis. Dried metabolites were resuspended in 100 ul 50% ACN:water and 5 ul was loaded onto a Luna NH2 3um 100A (150 × 2.0 mm) column (Phenomenex) using a Vanquish Flex UPLC (Thermo Scientific). The chromatographic separation was performed with mobile phases A (5 mM NH4AcO pH 9.9) and B (ACN) at a flow rate of 200 μl/min. A linear gradient from 15% A to 95% A over 18 min was followed by 7 min isocratic flow at 95% A and re-equilibration to 15% A. Metabolites were detected with a Thermo Scientific Q Exactive mass spectrometer run with polarity switching in full scan mode using a range of 70-975 m/z and 70.000 resolution. Maven (v 8.1.27.11) was used to quantify the targeted polar metabolites by AreaTop, using expected retention time and accurate mass measurements (< 5 ppm) for identification. Data analysis, including principal component analysis and heat map generation was performed using in-house R scripts. In brief, metabolite intensities (area under the curve) were normalized to protein content and analysed with one-way ANOVA (ANOVA column). Metabolites with a p-value < 0.05 were termed as significant. Heatmaps are z-score normalized assuming a normal distribution. Bar plots display relative amounts for each metabolite, calculated by averaging amounts for each condition (condition with the lowest average value set to 1).

### Metabolic assays

To measure metabolite levels in cells, cells were resuspended in an ice-cold homogenization medium (0.32 M sucrose, 1 mM EDTA, and 10 mM Tris–HCl, pH 7.4) and homogenized with 72 strokes using a tight pestle. After brief sonication at 40 % Amp, homogenates were centrifuged at 2000 g for 5 min at 4°C to remove the lipid layer and protein concentration was determined using BCA assay to normalize between conditions. 24 µg of protein was used per assay condition. Levels of xanthine + hypoxanthine (Abcam) in cell homogenates were measured using commercially available kits. Intracellular ATP was measured in living cells using ATP bioluminescence assay kit CLS II (Roche) according to the manufacturer’s instructions. Commercial kits were also used to measure xanthine + hypoxanthine (Abcam) directly in serum and conditioned medium. For more information, see the Key resources table.

### Quantification and statistical analysis

Experiments were conducted as 3 independent repeats or as indicated. Mice were randomly grouped in experimental groups and data were pooled from both sexes. Data were analysed and visualized using GraphPad Prism 9. The normal distribution of data was tested before applying parametric or nonparametric testing. For comparison between two independent groups, unpaired Student’s t-tests were applied. Comparisons across multiple groups were performed using one-way or two-way ANOVA with Šídák or Tukey multiple testing correction. Data were considered statistically significant when p value < 0.05.

### Data and code availability

RNA sequencing data reported in this paper is available at accession number GEO: GSE263837. Proteomics and metabolomics data can be accessed at XXXX.

