## Supplementary Information for "Autophagy acts as a brake on obesity-related fibrosis by controlling purine nucleoside signalling"

**Supplementary Figure 1: Validation of *Atg7^Ad^* mouse model to study the role of autophagy in diet-induced obesity.**

A) Relative mRNA expression of *Atg7* in gWAT of WT and *Atg7^Ad^* mice measured by qRT-PCR. n = 8-9 mice. Representative of 3 independent experiments.

B) Representative western blot analysis of LC-3I, LC-3II, and vinculin in gWAT to measure autophagy flux. WT and *Atg7^Ad^* mice were fed a normal chow diet (NCD) or high fat diet (HFD) for 16 weeks. The representative samples are shown of 3 independent experiments.

C) WT and *Atg7^Ad^* mice were fed a normal chow diet (NCD) or high fat diet (HFD) for 16 weeks before autophagy flux in gWAT was assessed as explained in Materials and Methods. Western blot analysis of autophagy flux (as in B) was calculated as (LC3‐II (Inh) – LC3‐II (Veh)). n = 3 – 6 mice. Representative of 3 independent experiments.

Data are presented as mean ± SD. Dots represent individual biological replicates. Statistical analysis by unpaired t-test (A) and one-way ANOVA with Tukey multi comparisons test (C).

**Supplementary Figure 2: Loss of autophagy induces a stress response and cell death, and profoundly disturbs mitochondrial homeostasis.**

A) Vulcano plot of cellular response to oxidative stress (GO:0034599).

B) Heatmap of OXPHOS biological process (GO:0006119). Proteins belonging to different mitochondrial complexes are colour-coded. Values were scaled by row (protein) using z-score.

C-D) Representative immunofluorescence staining of Perilipin-1 on gWAT sections from WT and *Atg7^Ad^* mice following HFD feeding for 16 weeks. (C) Quantification of Perilipin-1 staining as a percentage of Perilipin-1 negative adipocytes compared to total adipocytes from (C). Legend: blue = dapi, red = Perilipin-1. n = 7-8 mice. Data are merged from 3 independent experiments.

E) Cleaved caspase 3 was assessed by western blot analysis in gWAT explants cultured overnight *ex vivo* and treated with either DMSO or 20 µM Q-VD-OPh, a pan-caspase inhibitor. gWAT was isolated from WT and *Atg7^Ad^* mice fed with HFD for 16 weeks. n = 5-6 mice. Representative of 3 independent experiments.

F-G) Heatmap showing intracellular (F) and extracellular (G) adipokine accumulation. Lep = leptin, Adipoq = adiponectin. Values were scaled by row (protein) using z-score.

**Supplementary Figure 3: Metabolomics analysis reveals a prominent role of autophagy in amino acid metabolism.**

A) Principal component analysis (PCA) representation of metabolomics analysis. n = 3 mice per genotype, males only.

B) Relative abundance of uric acid measured with metabolomics analysis in adipocytes isolated from gWAT of WT and *Atg7^Ad^* mice following HFD feeding for 16 weeks. n = 3 mice.

C) Activity of xanthine oxidase (XO) after 2-hour incubation with WT and *Atg7^Ad^* adipocyte lysates. n = 11 mice. Data are merged from 3 independent experiments. Data are presented as mean ± SD. Dots represent individual biological replicates.

D) Z-score heatmap of significantly (p < 0.05) abundant metabolites in adipocytes. Metabolomics analysis was performed on adipocytes isolated from gWAT of WT and *Atg7^Ad^* mice following HFD feeding for 16 weeks. n = 3 mice.

E) Relative abundance of amino acids measured with metabolomics analysis in adipocytes isolated from gWAT of WT and *Atg7^Ad^* mice following HFD feeding for 16 weeks. n = 3 mice.

F) Log_2_ fold change heatmap of significantly differentially abundant proteins between WT and *Atg7^Ad^* involved in glycolysis, TCA cycle and FAO in adipocytes as measured by proteomics analysis.

All heatmap values were scaled by row (protein/metabolite) using z-score.

**Supplementary Figure 4: The absence of adipocyte autophagy ameliorates metabolic syndrome in diet-induced obese mice.**

WT and *Atg7^Ad^* mice were fed normal chow diet (NCD) or high fat diet (HFD) for 12-16 weeks prior to analysis.

A) Fat pad weight. n = 8-16 mice. Data are merged from 3 independent experiments.

B) Weight gain relative to starting weight over 12 weeks of NCD or HFD feeding. n = 4 mice. Representative of 3 independent experiments.

C) Caloric food intake between NCD and HFD-fed mice. n = 4 mice. Representative of 3 independent experiments.

D) Glucose tolerance test performed at 12 weeks of HFD feeding. n = 4 mice. Representative of 3 independent experiments.

E) Area under the curve (AUC) for glucose tolerance test for (E).

F) H&E staining of liver harvested from HFD-fed WT and *Atg7^Ad^* mice after 16 weeks of feeding. Examples of fat deposition marked with asterisk, scale bar, 200 µm. Representative of 3 biological replicates.

G) Quantification of white (lipid) area from G. n = 6-8 mice. Data are merged from 3 independent experiments.

H-K) Serum concentration of triglycerides (TG, H), high-density lipoprotein (HDL, I), adiponectin (J), and leptin (K). n = 3-20 mice. Data are merged from 3 independent experiments.

L) Quantification of adipocyte number and diameter in gWAT from HFD-fed mice. n = 6 mice. Representative of 3 biological replicates.

Data are presented as mean ± SD. Dots represent individual data points. Statistical analysis by two-way ANOVA with Tukey multi comparisons test (D, E), unpaired t-test (G), and one-way ANOVA (J-K).

**Supplementary Figure 5: Gating strategies to determine cellular composition of WAT in *Atg7^Ad^* mice.**

A) Gating strategy for CD45^+^, CD31^+^, and PDGFRα^+^ subsets within live cells.

B) Gating strategy for neutrophils (Ly6G^+^), eosinophils (Siglec-F^+^), macrophages (F4/80^+^ CD64^+^), natural killer T (NKT) cells (CD3^+^ NK1.1^+^), TCRβ^+^ CD8^+^ and CD4^+^ T cells within CD45^+^ live cells.
