## Supplementary Figures for "Autophagy acts as a brake on obesity-related fibrosis by controlling purine nucleoside signalling"

Sup. Fig. 1: Validation of *Atg7^Ad* mouse model to study the role of autophagy in diet-induced obesity.

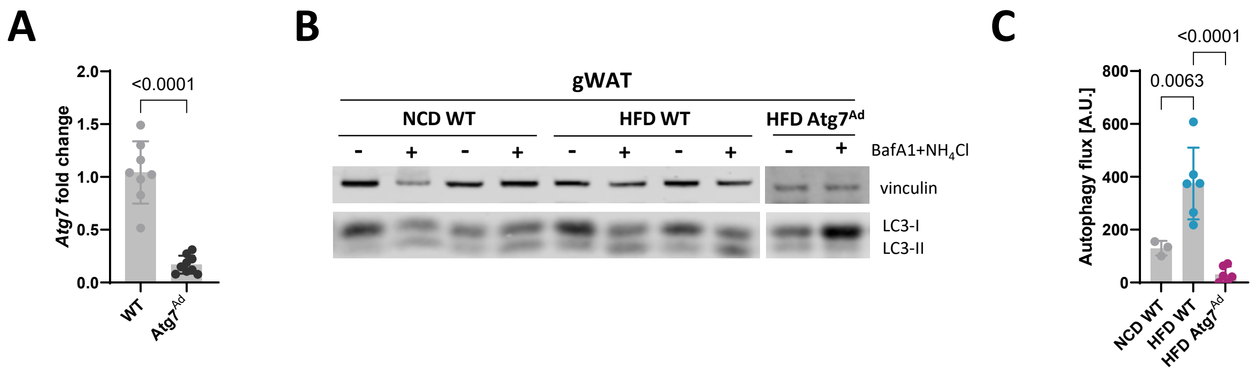

### Sup. Fig 2: Loss of autophagy induces a stress response and cell death, and profoundly disturbs mitochondrial homeostasis.

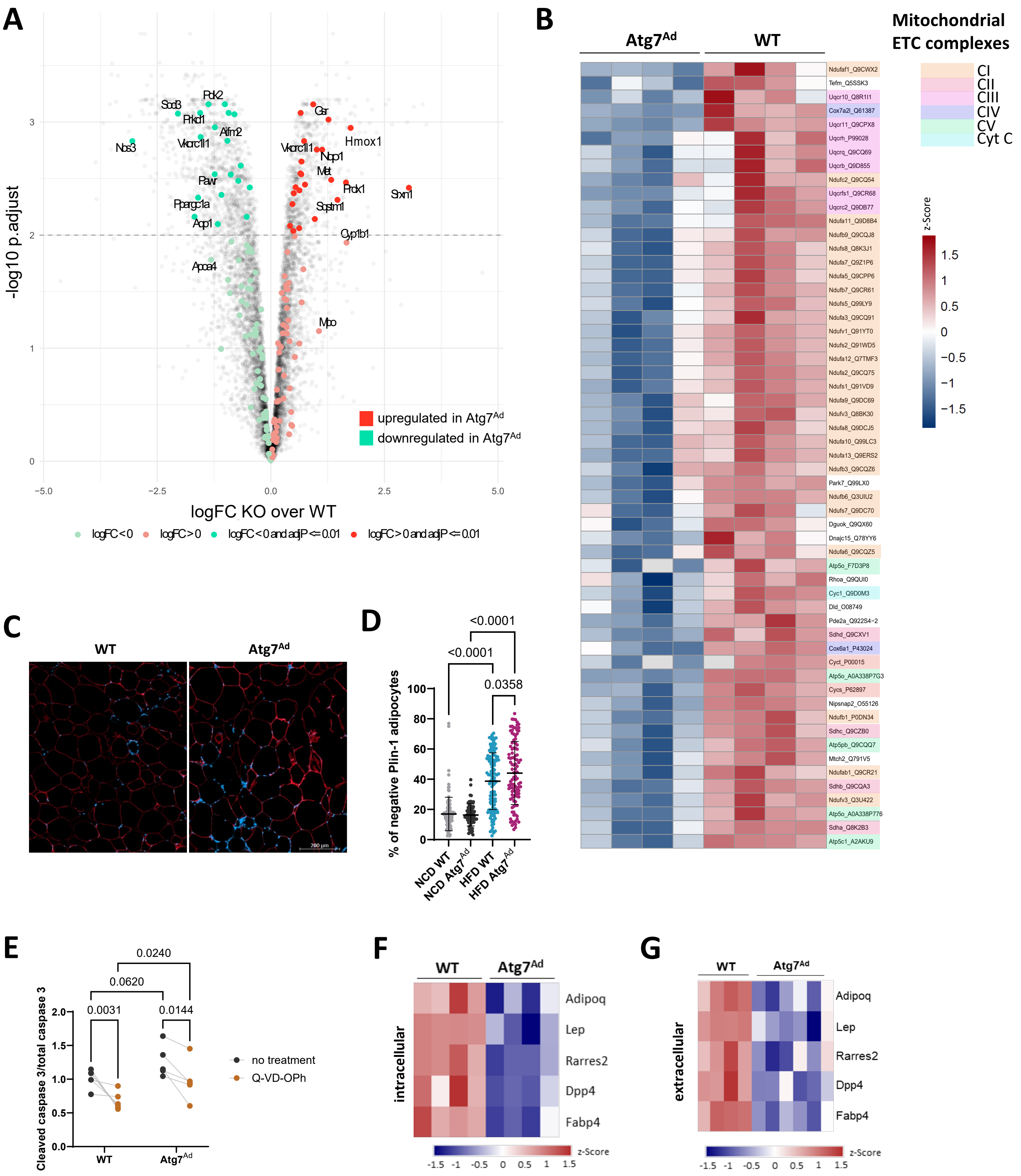

Sup. Fig 3: Metabolomics analysis reveals a prominent role of autophagy in amino acid metabolism.

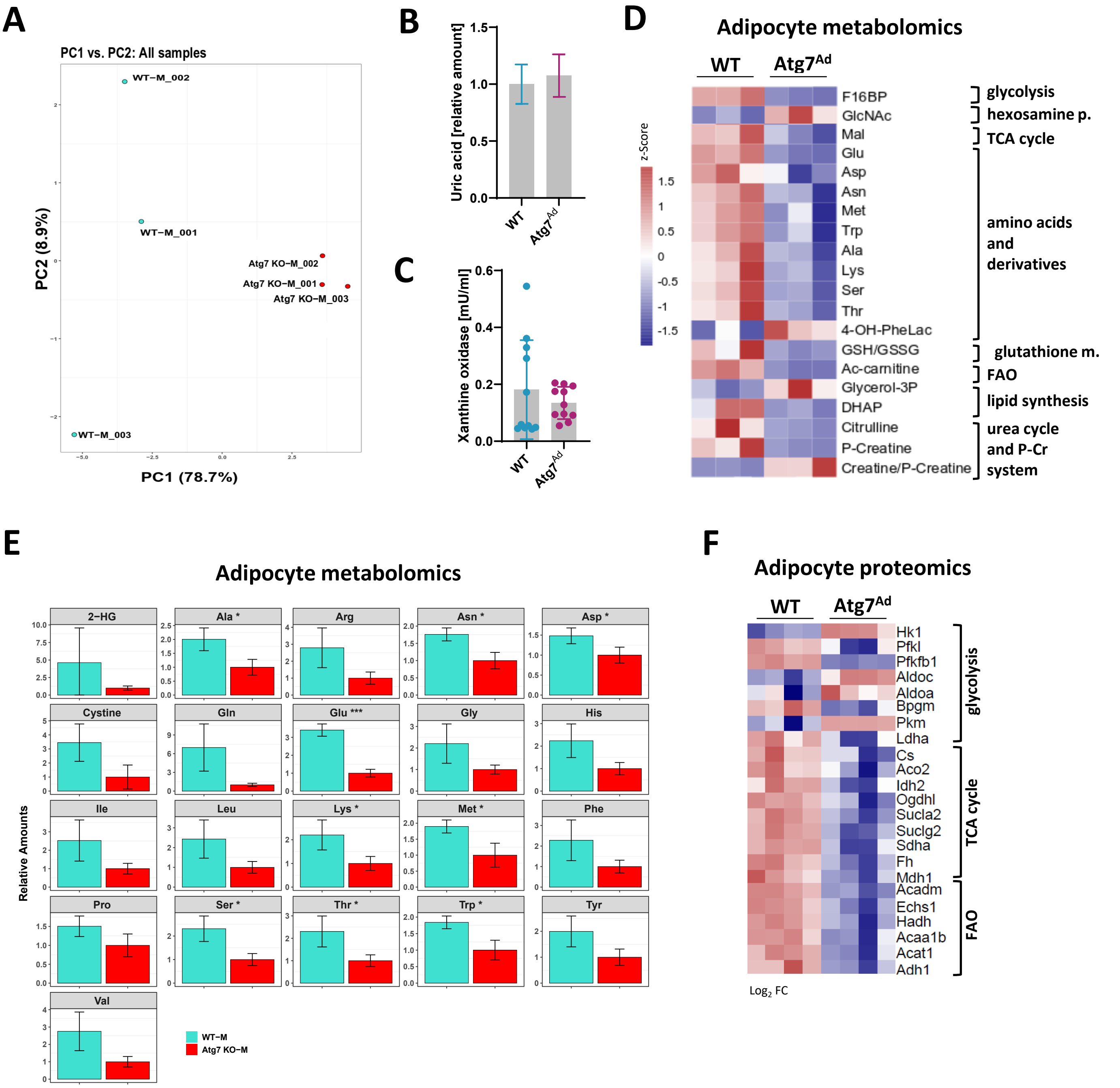

Sup. Fig 4: The absence of adipocyte autophagy ameliorates metabolic syndrome in diet-induced obese mice.

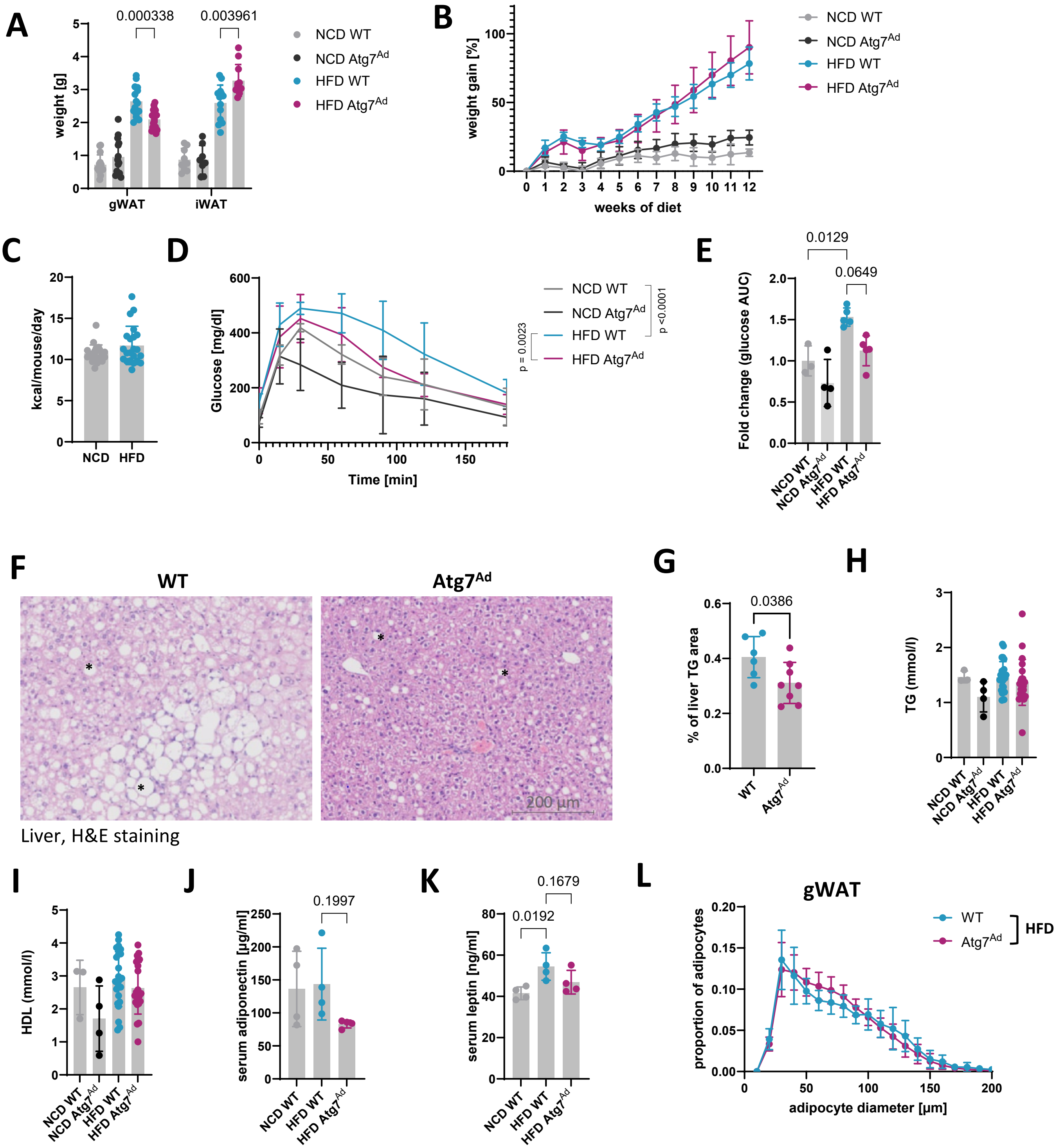

### Sup. Fig 5: Gating strategies to determine cellular composition of WAT in Atg7$^{Ad}$ mice.

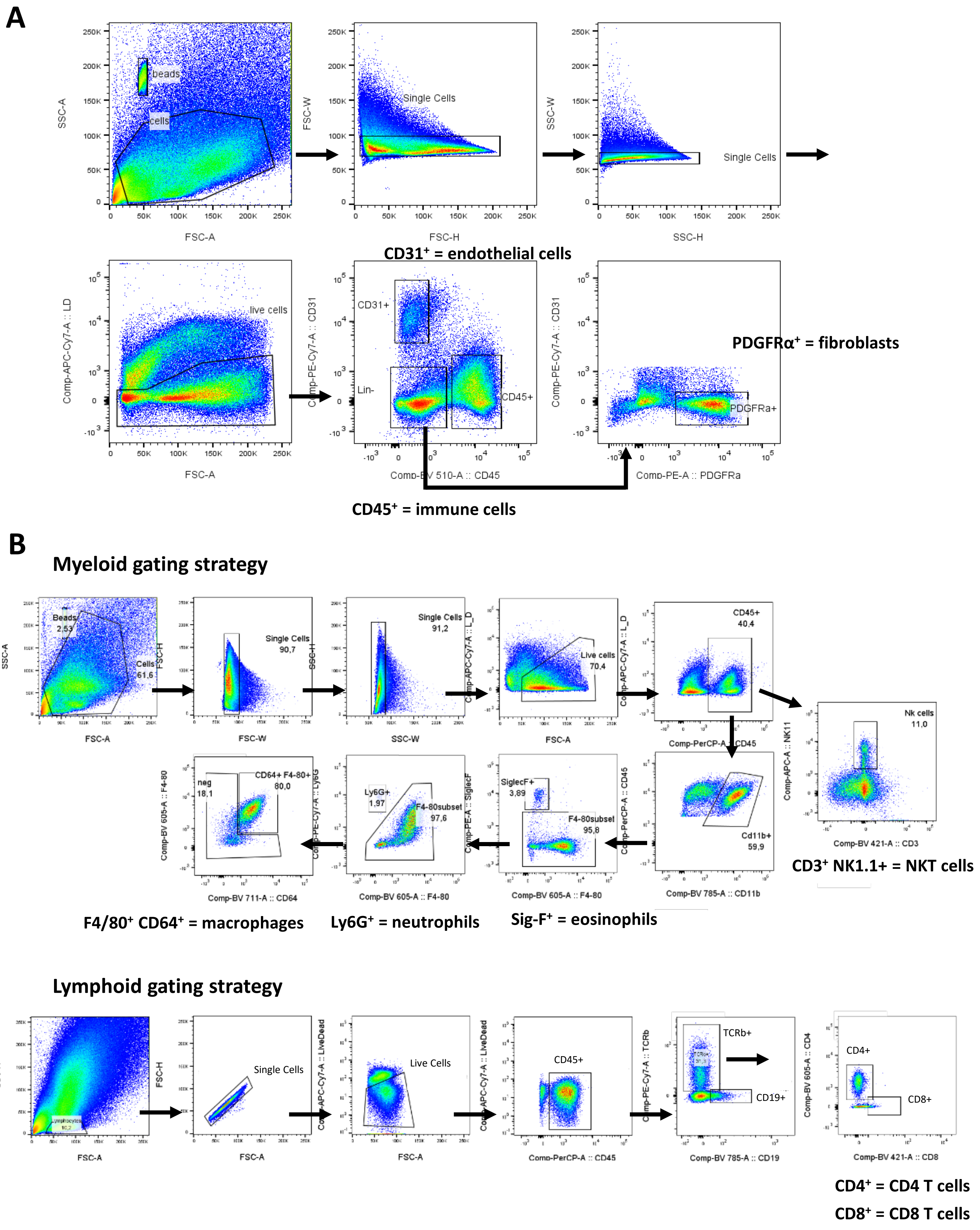
